# Bone Mechanosensing Dictates Hematopoietic Stem Cell Fate and Immune Homeostasis

**DOI:** 10.64898/2026.07.05.736555

**Authors:** Wuju Zhang, Yating Pan, Xiaoling Xie, Jianmeng Du, Huihui Zhang, Zhixin Ye, Xiaoqing Yan, Jionghong Huang, Hailin Jing, Sheng Zhang, Xiaolin Liu, Di Chen, Yongguang Liu, Xiao Yu, Xiaochun Bai

## Abstract

The vertebrate water-to-land transition was accompanied by a six-fold increase in gravitational force, followed by the migration of hematopoietic stem cells (HSCs) from kidney or liver to the bone marrow, and the acquisition of enhanced immune functions to cope with novel environmental pressures. Bone senses mechanical loading and provides a microenvironment for HSC development, yet whether bone mechanosensation affects immune cell development and immune homeostasis remains unclear. Here, we unveil bone as a mechanosensory organ that translates mechanical force into hematopoietic instructions. Mechanical loading of bone directs HSCs toward lymphoid lineages in mice and non-human primates, whereas unloading favors myeloid commitment. This process requires osteocyte mechanosensor Piezo1, which induces loading-responsive bone-derived factors such as IL1R2, SERPINC1 and INMT, thereby restraining inflammatory signaling and guiding HSC differentiation. Osteocyte Piezo1 deficiency recapitulates the hematopoietic and immune alterations observed during unloading. Functionally, this pathway enhances acute infection resistance and suppresses immunosenescence in mice and in aged long-tailed macaques. Notably, skeletal mechanoregulation of immune homeostasis is conserved across vertebrate species. Our research defines the mechano-bone-immune axis (mechano-osteoimmunology), offering a novel evolutionary perspective on the interconnected development of the skeletal and immune systems and presenting a promising non-pharmacological target to address immune dysfunction.

## Introduction

Hematopoietic stem cells sustain lifelong immune cell production, and their lineage decisions are tightly controlled by specialized bone marrow niches to maintain immune homeostasis. Although numerous cellular and molecular cues that govern HSC maintenance and differentiation have been identified,^1–3^ whether physiological signals arising from the organism as a whole are integrated into hematopoietic fate decisions remains poorly understood.

Mechanical loading generated by body weight and physical activity is a fundamental feature of vertebrate physiology.4-7 As the most abundant cells in bone, osteocytes account for 90% of bone cells and function as the principal mechanosensors of the skeleton, converting mechanical forces into biochemical signals that regulate bone remodeling and systemic physiology.8-12 Although osteoimmunology has revealed extensive reciprocal interactions between skeletal and immune systems,13,14 most studies have focused on how immune cells influence bone homeostasis, whereas whether skeletal mechanosensation actively regulates hematopoiesis and immunity remains largely unexplored. This question is particularly relevant given the longstanding association between physical activity and immune competence. Exercise has been linked to improved immune function and reduced chronic inflammation, whereas prolonged immobilization, bed rest, microgravity exposure, and aging-associated declines in mobility are frequently accompanied by immune dysfunction and altered hematopoiesis.15-17 Furthermore, during transition of vertebrates from water to land,14 the six-fold increase in gravitational force paralleled the relocation of the HSC niche to the bone marrow.18 These observations suggest that bone mechanical information may represent an important but poorly understood physiological input into immune regulation.

Piezo1 is a mechanically activated ion channel that mediates mechanotransduction in diverse tissues and cell types, including endothelial, epithelial, skeletal, and immune cells.19-21 In the skeleton, Piezo1 is required for osteocyte and osteoblast responses to mechanical loading and plays a central role in bone adaptation to force.22 However, whether osteocyte Piezo1 functions beyond skeletal homeostasis to relay mechanical information to the hematopoietic and immune system has not been reported.

Here, we identify a bone mechanosensory pathway that couples mechanical loading to HSC lineage allocation and systemic immune homeostasis. Mechanical loading of bone promotes commitment of HSC to the lymphoid lineage and inhibits immune aging, whereas diminished mechanosensing of bone favors myeloid differentiation and drives immunosenescence, a decision governed by the mechanosensor Piezo1 of osteocytes. Mechanistically, osteocytic Piezo1 drives the release of factors such as Interleukin-1 receptor type 2 (IL1R2), Serpin family C member 1 (SERPINC1) and Indolethylamine N-methyltransferase (INMT), to restrain IL-1 signaling and shape HSC fate decisions. Furthermore, activation of mechanical force-Piezo1 in osteocytes enhances immune competence, whereas its disruption compromises host defense and promotes immunosenescence. Importantly, this regulatory axis is conserved in non-human primates, where mechanical loading restores lymphopoiesis and mitigates hallmarks of immune aging in aged long-tailed macaques. Our findings establish osteocytes as critical intermediaries that communicate organismal mechanical status to the hematopoietic system and reveal a conserved mechanism linking bone mechanosensation to immune homeostasis across species, which could be manipulated to combat immune dysfunction from prolonged bed rest and spaceflight to decreased mobility during aging.

## Results

### Mechanical loading promotes lymphopoiesis, whereas unloading drives myeloid skewing

To assess the effects of mechanical force on hematopoiesis, a murine model of mechanical unloading was established using a four-week tail suspension (TS) protocol. Bone marrow cells from both TS and control mice were analyzed using single-cell RNA sequencing (scRNA-seq) to gain a comprehensive understanding of the changes within the hematopoietic compartment (Figure 1A). Unsupervised clustering of the scRNA-seq data identified all major hematopoietic cell types, including various mature and immature immune cells (Figure 1B and S1A–S1C). Comparative analysis revealed a marked shift in the composition of progenitor populations following mechanical unloading (Figure 1C and 1D). Specifically, the proportion of lymphoid-primed cells was decreased, while the proportion of myeloid-primed cells was increased in TS mice as compared to controls (Figure 1E). Trajectory analysis indicated that cells from the TS group were skewed toward myeloid differentiation (HSC→GMP), whereas control cells showed a greater propensity to progress along the lymphoid branch (HSC→CLP→ProB→PreB) (Figure 1F). Furthermore, the progenitor clusters were annotated using established gene signatures, and the cluster that was diminished under the unloading condition was enriched with genes associated with common lymphoid progenitors (CLPs), including *Tcf3* and *Il7r* (Figure 1G). Conversely, the expanding cluster expressed markers typical of granulocyte-macrophage progenitors (GMPs), such as *Csf2ra* and *Ly6c2* (Figure S1E). Additionally, myeloid-associated genes (*Hmgb2* and *Ifngr1*) were upregulated in the TS group, while lymphoid regulatory genes (*Ebf1*, *Ikzf1*, and *Bcl11a*) were downregulated (Figure S1E).

**Figure 1.**
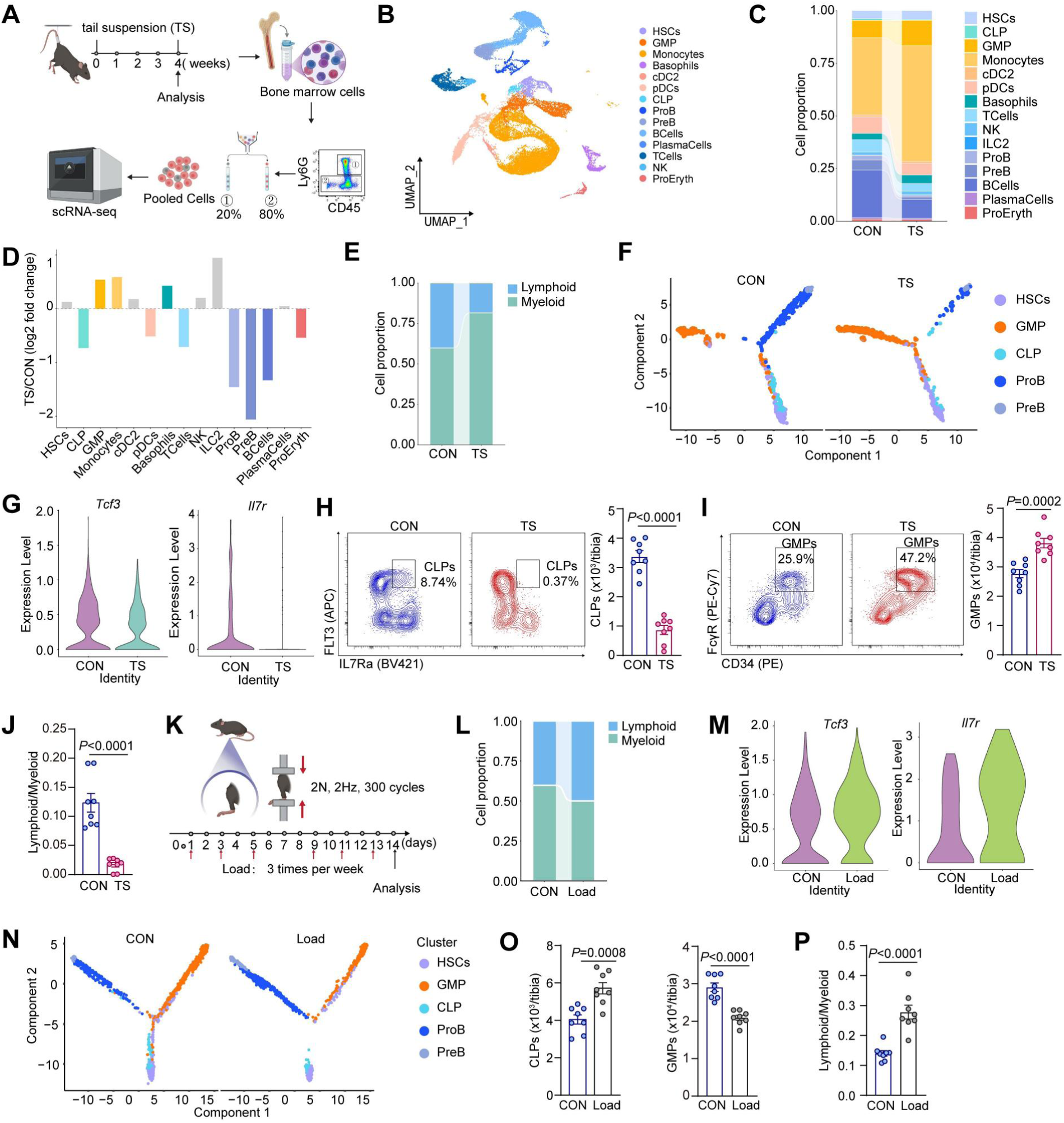
Mechanical unloading drives myeloid skewing, whereas dynamic tibial loading promotes lymphopoiesis in the bone marrow. **A,** Experimental design for mechanical unloading by tail suspension (TS; 4 weeks) and schematic of BM cell processing and scRNA-seq workflow. **B,** UMAP visualization of scRNA-seq profiles from BM hematopoietic cells, coloured by annotated cell types. **C,** Relative cell-type proportions in BM from control (CON) and TS mice based on scRNA-seq. **D,** Log₂ fold change of indicated cell-type frequencies in TS relative to CON mice. **E,** Quantification of lymphoid-versus myeloid-primed progenitor proportions in CON and TS BM. **F,** Trajectory (pseudotime) analysis of progenitor states showing TS cells biased toward the myeloid branch (HSC→GMP), whereas CON cells preferentially progress along the lymphoid branch (HSC→CLP→ProB→PreB). **G,** Violin plots showing reduced expression of lymphoid lineage–associated genes (*Tcf3* and *Il7r*) in progenitors from TS mice relative to controls. **H,** Representative flow cytometry plots and quantification of CLPs in CON and TS BM (*n* = 8). **I,** Representative flow cytometry plots of LSK progenitors showing GMP subsets, and quantification of GMPs in CON and TS mice (*n* = 8). **J,** Quantification of the lymphoid-to-myeloid output bias as the CLP/GMP ratio (*n* = 8). **K,** Experimental scheme for dynamic tibial axial loading (DTAL; 2 N, 2 Hz, 300 cycles per session, three sessions per week) followed by BM analysis. **L,** Quantification of lymphoid- versus myeloid-primed progenitor proportions in CON and DTAL (Load) mice. **M,** Violin plots showing increased *Tcf3* and *Il7r* expression in progenitors following DTAL. **N,** Pseudotime trajectory analysis of HSC/MPP differentiation illustrating preferential progression toward the lymphoid branch (HSC→CLP→ProB→PreB) after DTAL, compared with myeloid skewing (HSC→GMP) in controls. **O,** Flow cytometric quantification of CLPs and GMPs in CON and DTAL mice (*n* = 8). **P,** Ratio of lymphoid to myeloid progenitors in CON and DTAL mice (*n* = 8). For scRNA-seq analyses, *n* = 2 mice per group, with each biological replicate corresponding to one mouse. Neutrophils were removed prior to downstream scRNA-seq analyses. Data are presented as mean ± s.e.m. *P* values were calculated using two-sided unpaired Student’s *t*-test. *P* values are indicated in the plots.

To validate the scRNA-seq findings, flow cytometry of BM cells from both TS and control mice was conducted, utilizing a defined gating strategy to identify HSCs and progenitors (Figure S1F). Consistent with the scRNA-seq results, mechanical unloading significantly reduced the number of CLPs (Lin^-^Sca-1^lo^c-Kit^lo^IL-7Rα^+^FLT3^+^), while dramatically increasing the abundance of GMPs (Lin^-^Sca-1^-^c-Kit^+^CD34^+^FcγR^+^) within the BM (Figure 1H and 1I). Moreover, the ratio of CLPs to GMPs, an important indicator of lineage bias, was significantly lower for the TS mice, reflecting a systemic shift in hematopoietic output toward myeloid development (Figure 1J). Additionally, peripheral blood analysis showed increased abundances of neutrophils and Ly6C^+^ monocytes alongside reduced amounts of T and B cells (Figure S1G and S1H). Importantly, the frequency of upstream LSK cells (Lin-Sca-1^+^c-Kit^+^) remained stable, indicating that the observed effects pertained specifically to lineage-committed progenitors rather than the overall stem cell population (Figure S1I). Collectively, these results demonstrate that mechanical unloading significantly skews hematopoietic differentiation in the BM, prioritizing myeloid progenitor production over lymphoid progenitor development.

Having established that mechanical unloading enhances myeloid commitment, we sought to determine whether applying mechanical force could induce the opposite outcome. Dynamic tibial axial loading (DTAL) with a loading device is a precisely controlled method to apply a mechanical load specifically to bone.^23^ Mice were subjected to a moderate mechanical loading regimen using DTAL (Figure 1K). Following this intervention, BM cells were assessed via scRNA-seq and flow cytometry to evaluate changes to the hematopoietic progenitor compartment. The analysis revealed that mechanical loading effectively countered the effects induced by unloading. Specifically, the frequency of CLPs was significantly increased in mice subjected to DTAL as compared to controls, accompanied by corresponding decreased proportions of GMPs (Figure 1L). Expression analysis of lineage-defining genes further confirmed this trend, with a notable upregulation of *Tcf3* and *Il7r* in the progenitor cells of the loaded mice (Figure 1M). To further explore lineage commitment bias, a pseudotime analysis of the HSC and MPP populations was conducted. This trajectory analysis revealed that cells from the loaded group were more likely to follow a path leading to the lymphoid branch (HSC→CLP→ProB→PreB), while control cells showed a propensity toward myeloid differentiation (HSC→GMP) (Figure 1N). Flow cytometry provided clear quantitative evidence supporting the pro-lymphoid effects of mechanical loading. The proportion of CLPs was significantly elevated in the BM of the loaded mice, while the number of GMPs was markedly reduced (Figure 1O). Moreover, the ratio of CLPs to GMPs was significantly higher for the loaded mice (Figure 1P). These findings demonstrate that mechanical loading serves as a positive regulator of lymphoid development. When considered alongside the effects of unloading, these results establish a bidirectional regulatory role of mechanical forces in the fate of hematopoietic cells, where mechanical deprivation promotes myelopoiesis and mechanical stimulation supports lymphopoiesis.

### Osteocyte mechanosensing directs commitment of HSC

Osteocytes, which account for 90% of bone cells, are long-living, terminally differentiated cells and the principal mechano-sensors of the skeleton.^24,25^ Given that the skeleton is the primary organ exposed to gravitational and mechanical forces, we hypothesized that osteocytes act as key sensors that translate these forces into signals that influence hematopoietic differentiation. To investigate this premise, we analyzed the BM of osteocyte *Piezo1* knockout mice (*Dmp1*-Cre; *Piezo1*^fl/fl^, referred to as *Piezo1^Dmp^*^1^) alongside their control littermates (*Piezo1*^fl/fl^) under standard living conditions (Figure S2A). Single-cell transcriptional profiling of the hematopoietic compartment revealed apparent remodeling of progenitor composition of progenitor population in *Piezo1^Dmp^*^1^ mice as compared to controls (Figure 2A–2C). Even in the absence of mechanical intervention, the proportion of lymphoid-primed progenitors was decreased, while the abundance of myeloid-primed progenitors was increased in the *Piezo1^Dmp^*^1^ cohort (Figure 2A–2C). Trajectory analysis indicated that cells from the *Piezo1^Dmp^*^1^ mice were skewed toward myeloid differentiation (HSC→GMP), whereas cells from CON showed a greater propensity to progress along the lymphoid branch (HSC→CLP→ProB→PreB) (Figure 2D). Further examination through differential gene expression and pathway analyses indicated that the HSC/MPP population of *Piezo1^Dmp^*1 mice exhibited upregulation of myeloid-associated gene signatures alongside a downregulation of genes involved in lymphoid lineage commitment (Figure 2E). Flow cytometry confirmed these transcriptional observations, showing significantly reduced CLP numbers and markedly increased GMP numbers in the BM of *Piezo1^Dmp^*^1^ mice as compared to controls (Figure 2F–2H). Notably, the total number of LSK cells remained unchanged, indicating that the absence of osteocytic *Piezo1* specifically affects lineage commitment rather than the overall populations of stem and progenitor cells (Figure S2B). These findings demonstrate that the loss of the mechanosensor *Piezo1* in osteocytes leads to a shift of non-autonomous in hematopoiesis cells toward the myeloid lineage under homeostatic conditions. This phenotype closely resembles the effects observed with systemic mechanical unloading, supporting the notion that osteocytes play a crucial role as mechanosensory cells that regulate the balance between lymphoid and myeloid differentiation in the BM niche through *Piezo1*-dependent signaling. To further investigate the role of osteocyte-derived *Piezo1* in regulation of hematopoietic fate, DTAL was applied to *Piezo1^Dmp^*^1^ mice alongside their control littermates (Figure 2I). Flow cytometry revealed that in *Piezo1*^f/f^ control mice, DTAL significantly enhanced the frequency of CLPs while decreasing that of GMPs, thereby confirming our earlier observations (Figure 2J). However, in *Piezo1^Dmp^*^1^ mice, DTAL had no effect on the frequencies of either CLPs or GMPs (Figure 2J). As a result, the loading-induced increase in the CLP/GMP ratio was evident solely in control mice (Figure 2K). These findings underscore the critical importance of osteocytic *Piezo1* in mediating the impact of mechanical loading on lymphoid lineage commitment.

**Figure 2.**
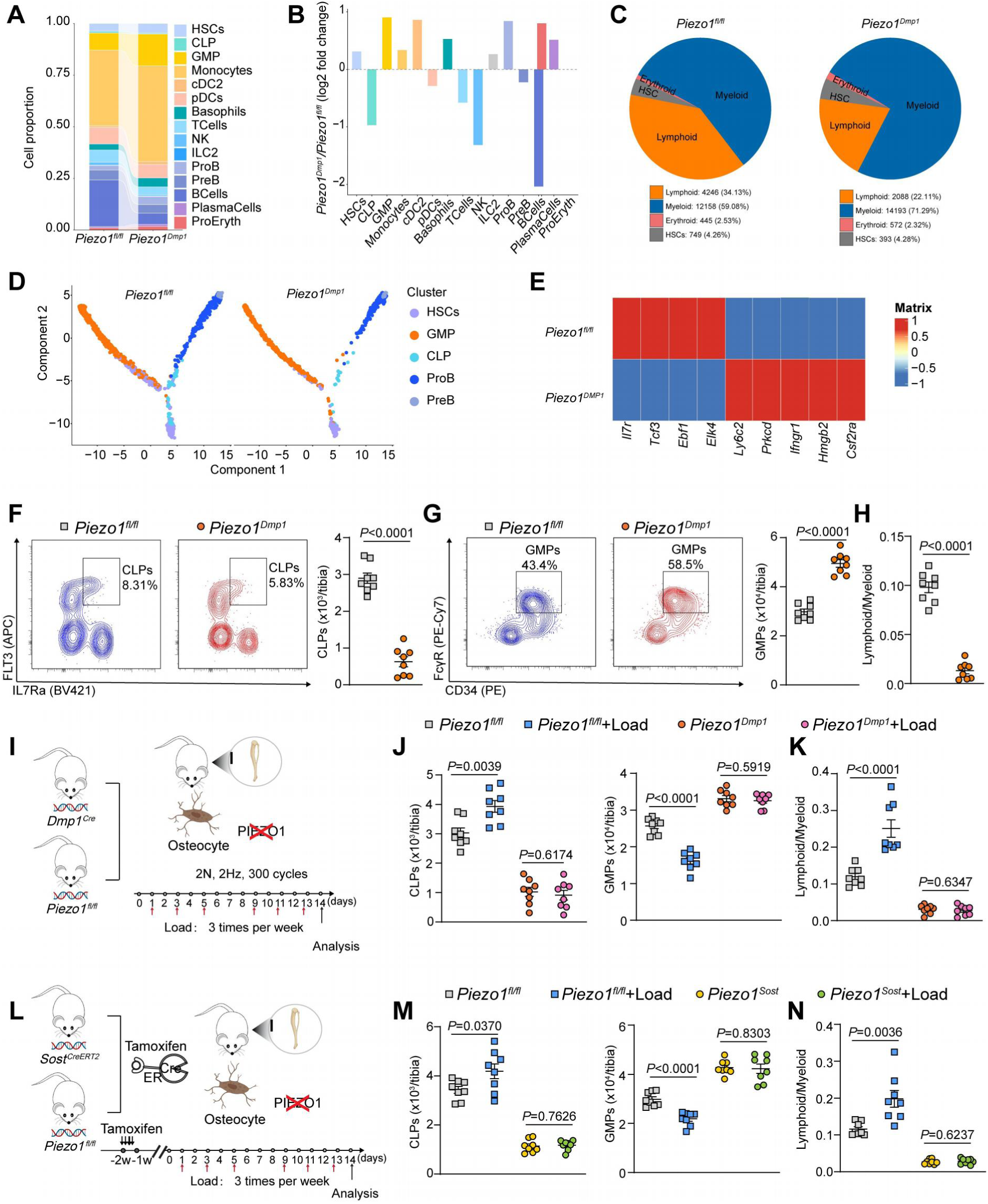
Osteocytic Piezo1 is required for mechanical force to promote lymphoid commitment of haematopoietic progenitors. **A,** Stacked bar plot showing the proportions of major BM hematopoietic populations in *Piezo1*^fl/fl^ and *Piezo1*^Dmp1^, as determined by scRNA-seq. **B,** Log_2_ fold change in the frequency of indicated BM hematopoietic populations in *Piezo1^Dmp1^* versus *Piezo1*^fl/fl^ mice. **C,** Lineage distribution (lymphoid, myeloid, erythroid and HSC compartments) derived from scRNA-seq in *Piezo1*^fl/fl^ and *Piezo1^Dmp1^* mice. **D,** Pseudotime trajectory analysis of HSC/MPP differentiation showing skewing toward the myeloid branch (HSC→GMP) in *Piezo1^Dmp1^*mice compared with preferential progression toward the lymphoid branch (HSC→CLP→ProB→PreB) in controls. **E,** Heat map of lineage-associated gene signatures in progenitor populations from *Piezo1*^fl/fl^ and *Piezo1^Dmp1^* mice, showing reduced lymphoid regulatory genes (*Il7r*, *Tcf3*, *Ebf1*) and increased myeloid-associated genes (*Ly6c2*, *Prkcd*, *Ccr2*, *Hmgb2*, *Csf2ra*) in *Piezo1^Dmp1^* mice. **F,** Representative flow cytometry plots and quantification of CLPs in *Piezo1*^fl/fl^ and *Piezo1^Dmp1^*mice (*n* = 8). **G,** Representative flow-cytometry plots and quantification of GMPs in BM from *Piezo1*^fl/fl^ and *Piezo1^Dmp1^*mice (*n* = 8). **H,** Ratio of lymphoid to myeloid progenitors (CLP/GMP) in *Piezo1*^fl/fl^ and *Piezo1^Dmp1^* mice (*n* = 8). **I,** Experimental scheme for DTAL in *Piezo1*^fl/fl^ and *Piezo1^Dmp1^* mice. Mechanical loading was applied at 2 N, 2 Hz, 300 cycles, three times per week, followed by bone marrow analysis. **J,** Flow-cytometry quantification of BM CLPs and GMPs following DTAL in *Piezo1*^fl/fl^ and *Piezo1^Dmp1^*mice (*n* = 8). **K,** CLP/GMP ratio following DTAL in *Piezo1*^fl/fl^ and *Piezo1^Dmp1^* mice (*n* = 8). **L,** Inducible osteocyte-specific *Piezo1* deletion strategy with tamoxifen administration in adult mice, followed by DTAL. **M,** Flow-cytometry quantification of BM CLPs and GMPs in *Piezo1*^fl/fl^ and *Piezo1^Sost^* mice with or without DTAL (*n* = 8). **N,** CLP/GMP ratio in *Piezo1*^fl/fl^ and *Piezo1^Sost^* mice with or without DTAL (*n* = 8). For scRNA-seq analyses in **A**–**E**, *n* = 2 mice per group, with each biological replicate corresponding to one mouse. For flow cytometry analyses in **F**–**H** and **J**–**N**, *n* = 8 mice per group; each dot represents one mouse. Data are presented as mean ± s.e.m. For two-group comparisons, *P* values were calculated using two-sided unpaired Student’s *t*-test. For experiments involving genotype and mechanical loading, statistical significance was assessed using two-way ANOVA followed by multiple-comparisons correction. Exact *P* values are indicated in the plots.

To eliminate concerns of developmental compensation inherent in the constitutive *Dmp1*-Cre model, an inducible osteocyte-specific knockout model (*Sost-*CreERT2; *Piezo1*^fl/fl^) was adopted, referred to as *Piezo1^Sost^*(Figure 2L). In this approach, tamoxifen was administered to adult mice to selectively delete *Piezo1* in osteocytes (Figure S2C) prior to mechanical loading. Subsequent flow cytometry of the BM reaffirmed the necessity of osteocytic *Piezo1* in adult mice to respond to mechanical stress. Consistent with previous results, mechanical loading significantly elevated the frequency of CLPs while decreasing GMPs in the control mice (Figure 2, L and M). In contrast, this mechanotransductive response was entirely absent in the adult-induced *Piezo1^Sost^* mice (Figure 2M). Following tamoxifen-driven deletion of *Piezo1*, the hematopoietic progenitor pool of these mice exhibited no significant changes to the frequencies of CLPs or GMPs relative to the controls (Figure 2M). The CLP/GMP ratio rose substantially in response to mechanical loading in the control group but remained unchanged in *Piezo1^Sost^*mice (Figure 2N). These results compellingly demonstrate that the mechanosensory function of *Piezo1* in osteocytes is crucial to translate mechanical signals that govern hematopoietic lineage commitment, particularly by maintaining the balance between lymphoid and myeloid differentiation within the adult BM niche.

### Mechanical force induces bone-derived factors to skew hematopoiesis toward the lymphoid lineage

Having identified *Piezo1* in osteocytes as a pivotal regulator of hematopoietic lineage determination, we hypothesized that mechanical loading enhances the secretion of specific bone-derived factors that promote lymphoid lineage commitment.^26^ To this end, quantitative proteomic analysis of BM supernatants from control and DTAL mice was conducted. This analysis revealed a set of candidate proteins with significantly increased secretion following mechanical loading, including IL1R2, INMT, SERPINA3M, C9, SERPINC1 and TRIM21 (Figure 3A; S3A and S3B). Notably, this loading-induced protein signature contained multiple factors with potential immunomodulatory functions: IL1R2 acts as a soluble decoy receptor that limits IL-1 signaling^27,28^, SERPINC1 has been implicated in restraining coagulation-associated inflammation and endothelial activation^29^, and INMT is linked to indoleamine metabolism with reported anti-inflammatory and immune-regulatory potential^30^. Further quantitative PCR analysis of isolated osteocytes confirmed that several of these proteins, particularly IL1R2, SERPINC1 and INMT, were transcriptionally upregulated in response to mechanical stimulation (Figure 3B). To determine whether the loading-induced increase in IL1R2 was dependent on the mechanosensory function of osteocytes, IL1R2 levels were measured in *Piezo1^Sost^* mice. As predicted, mechanical loading led to a significant rise in IL1R2 protein levels in the BM of control mice. In contrast, *Piezo1^Sost^*mice displayed markedly lower baseline levels of IL1R2, and mechanical loading did not elevate IL1R2 levels in these mice (Figure 3C and 3D; S3C and S3D). These findings indicate that IL1R2 may serve as a crucial mechano-responsive factors, as upregulation post-loading was reliant on *Piezo1* activation in osteocytes, thereby positioning IL1R2 as a key mediator of mechanical signals from bone to the hematopoietic system.

**Figure 3.**
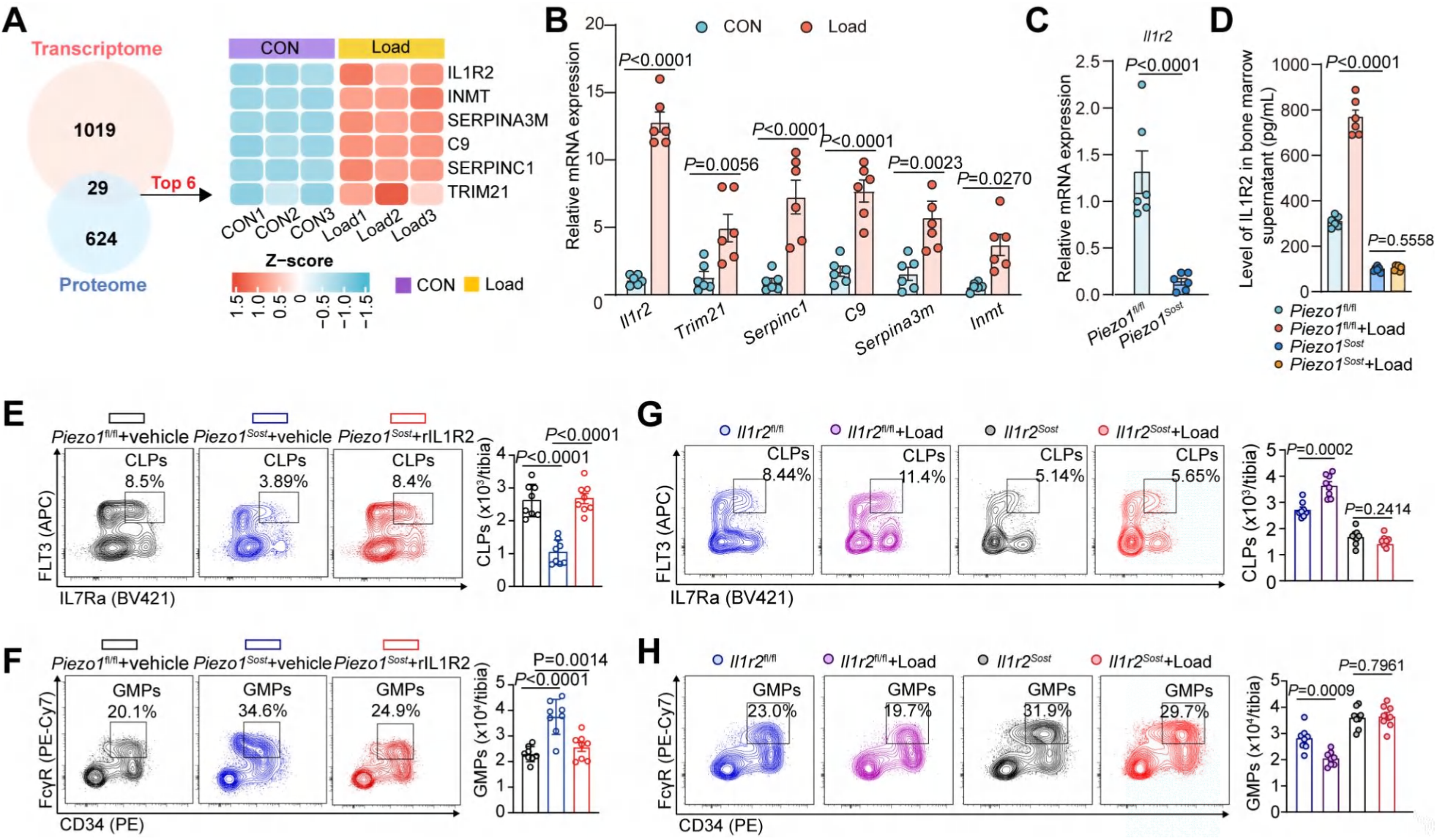
Mechanical loading induces osteocyte-derived IL1R2 to promote lymphoid commitment. **A,** Integration of bone transcriptome and BM supernatant proteomics identifies mechano-responsive osteokines increased after DTAL, highlighting IL1R2, INMT, C9, SERPINA3M, SERPINC1 and TRIM21; heat map shows z-scored expression/abundance across biological replicates. **B,** Quantitative PCR of isolated osteocytes showing loading-induced upregulation of candidate osteokine transcripts, including *Il1r2*, *Trim21*, *Serpinc1*, *C9*, *Serpina3m* and *Inmt* (*n* = 6). **C,** Relative *Il1r2* mRNA expression in osteocytes from control and *Piezo1^Sost^* (*n* = 6). **D,** Concentration of soluble IL1R2 protein in BM supernatants from *Piezo1*^fl/fl^ and *Piezo1^Sost^* mice with or without DTAL (*n* = 6). **E, F,** Representative flow-cytometry plots and quantification of CLPs **(E)** and GMPs **(F)** in *Piezo1*^fl/fl^ and *Piezo1*^Sost^ mice treated with vehicle or recombinant soluble IL1R2 (*n* = 8). **G, H,** Representative flow-cytometry plots and quantification of CLPs **(G)** and GMPs **(H)** in *Il1r2*^fl/fl^ and *Il1r2^Sost^* with or without DTAL (*n* = 8). Data are mean ± s.e.m.; each dot represents one mouse. For two-group comparisons, *P* values were calculated using two-sided unpaired Student’s *t*-test. For comparisons involving more than two groups, one-way ANOVA or two-way ANOVA followed by multiple-comparisons correction was used as appropriate. Exact *P* values are indicated in the plots.

To establish a direct causal link between IL1R2 and the regulation of hematopoiesis by mechanotransduction, we investigated whether administration of recombinant soluble IL1R2 could compensate for the mechanosensing impairment of *Piezo1^Sost^* mice. As previously shown, the proportion of CLPs was significantly reduced in *Piezo1^Sost^*mice as compared to controls. Remarkably, treatment with recombinant soluble IL1R2 significantly increased the CLP population in *Piezo1^Sost^*mice, while concurrently decreasing the GMP population, restored the CLP and GMP compartments toward control levels (Figure 3E and 3F). To further investigate the necessity of IL1R2 in the pro-lymphoid shift induced by mechanical loading, we generated osteocyte-specific *Il1r2*-deficient mice (*Sost-*CreERT2; *Il1r2*^fl/fl^, referred to as *Il1r2^Sost^*) and subjected them to DTAL (Figure S3E). In control mice, mechanical loading elicited the expected increase in CLPs accompanied by a decrease in GMPs. In contrast, this pro-lymphoid shift was completely abolished in *Il1r2^Sost^*mice (Figure 3G and 3H). These results underscore the role of IL1R2 as a critical downstream effector in the *Piezo1*-dependent signaling pathway linking osteocytes to hematopoietic progenitors, highlighting its significance in mechanoregulation of hematopoiesis.

To fully elucidate the molecular pathway that connects mechanical sensing to hematopoietic fate, we examined the signaling cascade downstream of PIEZO1 in osteocytes. PIEZO1 activation rapidly triggered NF-κB signaling and enhanced expression of IL1R2 in osteocytes, as indicated by increased phosphorylation of NF-κB p65 following Yoda1 stimulation, whereas pharmacological inhibition of PIEZO1 with GsMTx4 attenuated this response (Figure S3F and S3G). Moreover, blockade of NF-κB with Bay 11-7085 abolished Yoda1-induced IL1R2 upregulation, supporting a PIEZO1-NF-κB axis upstream of IL1R2 induction in osteocytes (Figure S3H). Inhibition of PIEZO1 using GsMTx4 largely abolished restoration of CLP pools following DTAL, as well as the shift from the myeloid to lymphoid lineage (Figure S3I and 3J). Together, these findings suggest that mechanical stimuli activate the Piezo1-NF-κB signaling pathway in osteocytes, which promotes upregulation of IL1R2. This osteokine inhibits IL-1 signaling, thereby creating a de-inflammatory microenvironment that reprograms hematopoiesis towards a lymphoid output and rejuvenates BM function.

### Bone mechanical cues regulate systemic inflammatory resilience during septic challenge

To investigate the functional implications of the mechano-regulated shift in hematopoiesis, mice were challenged with systemic inflammatory insults. Sepsis in TS and control mice was modeled using cecal ligation and puncture (CLP surgery) as well as lipopolysaccharide (LPS) injection.^31^ Following these challenges, TS mice exhibited a hyper-inflammatory response, with significantly elevated serum levels of IL-6, tumor necrosis factor (TNF)-α, and IL-1β compared to controls (Figure 4A and S4A). This systemic cytokine storm was accompanied by increased lung inflammation and, importantly, a markedly reduced survival rate of TS mice (Figure 4B–4D and S4B–4D). The administration of recombinant soluble IL1R2 to TS mice resulted in significantly improved survival, highlighting the role of IL1R2 in mediating susceptibility to injury (Figure 4E–4H and S4E–S4H). In contrast, mechanical loading conferred a notable protective advantage. Mice subjected to DTAL prior to sepsis induction demonstrated lower systemic inflammation and significantly improved survival rates as compared to their unloaded counterparts (Figure 4I–4K).

**Figure 4.**
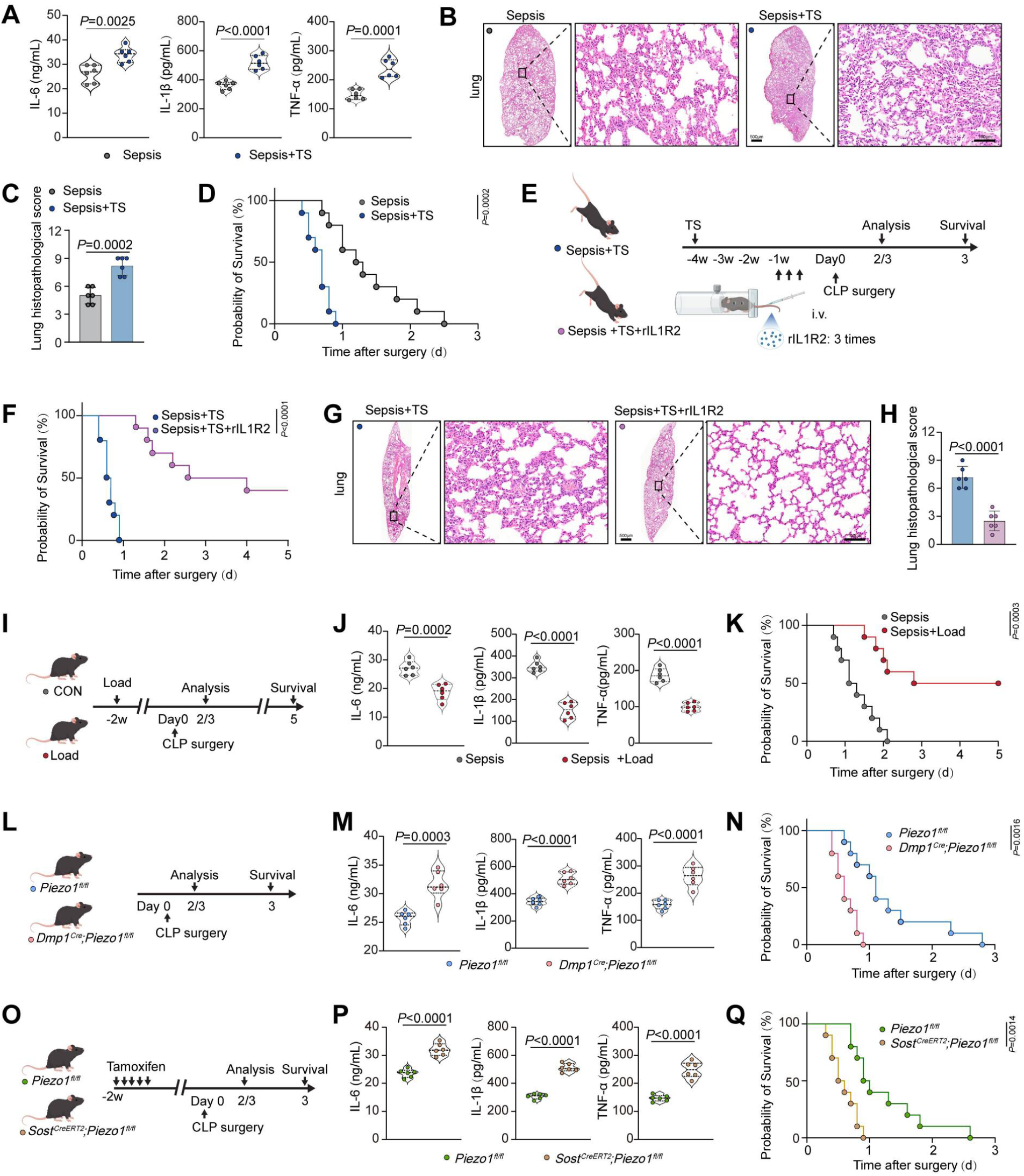
Mechanical cues in bone govern systemic inflammatory tolerance and survival during septic challenge. **A,** Serum cytokine concentrations of IL-6, IL-1β and TNF-α in mice subjected to cecal ligation and puncture surgery with or without prior tail suspension (*n* = 6). **B,** Representative H&E-stained lung sections from septic mice with or without prior tail suspension. Boxed regions are shown at higher magnification. Scale bars, 500 μm for overview images and 100 μm for magnified images (*n* = 6). **C,** Quantification of lung histopathology scores in septic mice with or without prior tail suspension (*n* = 6). **D,** Kaplan–Meier survival curves following cecal ligation and puncture surgery in mice with or without prior tail suspension (*n* = 10). **E,** Experimental scheme for recombinant soluble IL1R2 administration in tail-suspended mice undergoing cecal ligation and puncture surgery. **F,** Kaplan–Meier survival curves of tail-suspended septic mice treated with vehicle or recombinant soluble IL1R2. **G,** Representative H&E-stained lung sections from tail-suspended septic mice treated with vehicle or recombinant soluble IL1R2. Boxed regions are shown at higher magnification. Scale bars, 500 μm for overview images and 100 μm for magnified images (*n* = 6). **H**, Quantification of lung histopathology scores in tail-suspended septic mice treated with vehicle or recombinant soluble IL1R2 (*n* = 6). **I,** Experimental scheme for dynamic tibial axial loading before cecal ligation and puncture surgery. **J,** Serum concentrations of IL-6, IL-1β and TNF-α following cecal ligation and puncture surgery in non-loaded and mechanically loaded mice (*n* = 6). **K,** Kaplan–Meier survival curves following cecal ligation and puncture surgery in non-loaded and mechanically loaded mice (*n* = 10). **L,** Experimental scheme for septic challenge in *Piezo1*^fl/fl^ and *Piezo1^Dmp1^*. **M,** Serum cytokine concentrations (IL-6, IL-1β and TNF-α) following cecal ligation and puncture surgery in *Piezo1*^fl/fl^ and *Piezo1^Dmp1^*(*n* = 6). **N,** Kaplan–Meier survival curves following cecal ligation and puncture surgery in *Piezo1*^fl/fl^ and *Piezo1^Dmp1^*mice (*n* = 10). **O,** Experimental scheme for septic challenge in *Piezo1^Sost^* and controls after tamoxifen induction. **P,** Serum cytokine concentrations (IL-6, IL-1β and TNF-α) following cecal ligation and puncture surgery in *Piezo1*^fl/fl^ and *Piezo1^Sost^* mice (*n* = 6). **Q,** Kaplan–Meier survival curves following cecal ligation and puncture surgery in *Piezo1*^fl/fl^ and *Piezo1^Sost^*mice (*n* = 10). Data are mean ± s.e.m.; each dot represents one mouse. *P* values are shown in the panels; for cytokine measurements, two-sided unpaired Student’s *t*-test was used for two-group comparisons. Lung histopathology scores were compared using the Mann–Whitney U test. Survival differences were assessed using the log-rank (Mantel–Cox) test.

To further confirm the role of osteocyte mechanosensing in immune defense, we utilized constitutive (*Piezo1^Dmp^*^1^) and inducible (*Piezo1^Sost^*) osteocyte-specific knockout mice. Both *Piezo1^Dmp1^* and *Piezo1^Sost^* mice exhibited the same susceptible phenotype as TS mice, characterized by heightened production of inflammatory cytokines and decreased survival following septic challenge (Figure 4L–4Q). These results underscore the importance of the mechano-bone-immune axis as a critical regulator of systemic immune competence. Disruption of this axis, whether through environmental factors like unloading or genetic alterations such as Piezo1 deletion, renders the host more vulnerable to severe inflammation and mortality. Conversely, enhancing this pathway through mechanical loading appears to bolster host defenses and promote homeostasis.

### Mechanical loading alleviates age-related systemic inflammation and immunosenescence

Having established the significance in acute immune responses, we next examined the impact of mechanical sensing on chronic immune homeostasis during aging. Given the correlation between myeloid skewing (characterized by an elevated GMP/CLP ratio) and immunosenescence^32,33^, we investigated whether the mechano-bone-immune axis influences age-related immune decline. Subjecting aged WT mice to long-term (4 weeks) DTAL reversed age-related myeloid skewing (Figure 5A and 5B). Relative to young mice, aged mice displayed a canonical immunosenescent phenotype in the BM and peripheral blood, with reduced populations of B and T cells and expansion of Ly6C^hi^ monocytes, neutrophils, and CD11b^+^ myeloid cells. DTAL significantly mitigated these changes, increasing lymphoid compartments and suppressing myeloid expansion in both the BM and peripheral blood (Figure S5A–S5D). Meanwhile, aging led to marked accumulation of age-associated B cells in the peripheral blood and spleen, which was significantly reduced by DTAL (Figure 5C and 5D). In parallel, DTAL increased the frequency of CD44^lo^CD62L^hi^ naive T cells, in both the peripheral blood and spleen, counteracting the age-driven contraction of the naive T cell pool (Figure 5E and 5F; S5E and S5F). At the progenitor level, aging was associated with reduced CLPs and an imbalanced lymphoid-to-myeloid output. DTAL robustly restored the abundance of CLPs, improved the lymphoid/myeloid ratio, and reduced the age-associated expansion of GMPs (Figure 5G and 5H). Consistently, serum IL-6 and TNF-α levels were elevated in aged mice but significantly reduced by DTAL, indicating attenuation of inflammaging (Figure 5I).

**Figure 5.**
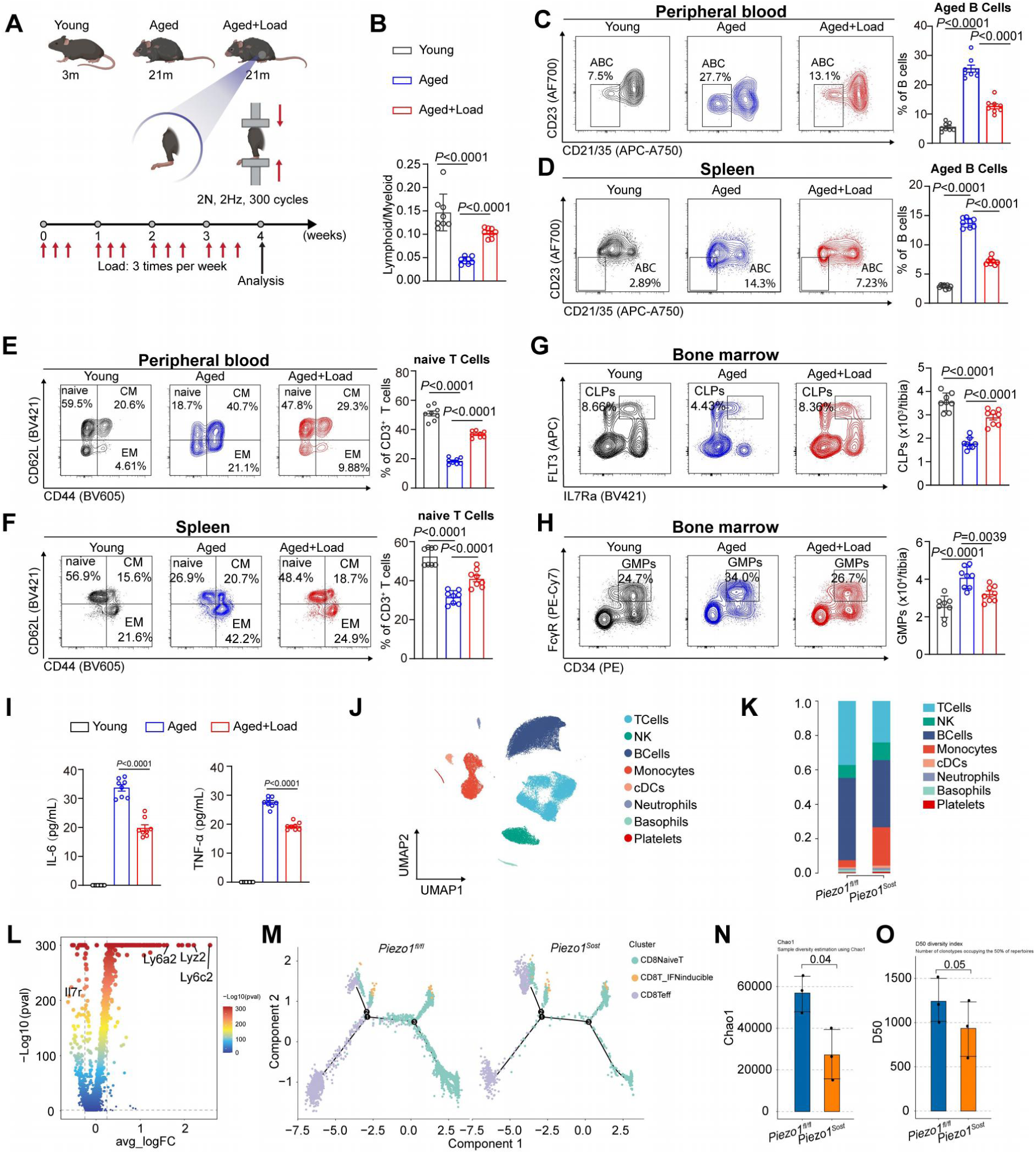
Mechanical bone loading mitigates age-associated systemic inflammation and immunosenescence via osteocyte mechanosensing. **A,** Experimental scheme of the long-term DTAL regimen applied to young and aged wild-type mice. **B,** Lymphoid-to-myeloid ratio in bone marrow from young, aged and aged+Load mice. **C,** Representative flow cytometry plots and quantification of age-associated B cells in peripheral blood from young, aged and aged+Load mice. Age-associated B cells were gated as CD23⁻CD21/35⁺ B cells. **D,** Representative flow cytometry plots and quantification of age-associated B cells in spleen from young, aged and aged+Load mice. **E,** Representative flow cytometry plots and quantification of CD44^lo^CD62L^hi^ naive T cells among CD3⁺ T cells in peripheral blood from young, aged and aged+Load mice. Central memory-like and effector memory-like T cell populations are also indicated. **F,** Representative flow cytometry plots and quantification of CD44^lo^CD62L^hi^ naive T cells among CD3⁺ T cells in spleen from young, aged and aged+Load mice. **G,** Representative flow cytometry plots and quantification of CLP numbers in bone marrow from young, aged and aged+Load mice. **H,** Representative flow cytometry plots and quantification of GMP numbers in bone marrow from young, aged and aged+Load mice. **I,** Serum IL-6 and TNF-α levels in young, aged and aged+Load mice. **J,** UMAP visualization of PBMC scRNA-seq profiles from *Piezo1*^fl/fl^ and *Piezo1^Sost^* mice, coloured by annotated immune cell type. **K,** Relative proportions of major PBMC immune populations in *Piezo1*^fl/fl^ and *Piezo1^Sost^*mice based on scRNA-seq. **L,** Volcano plot showing differentially expressed genes in PBMCs from *Piezo1^Sost^* mice compared with *Piezo1*^fl/fl^ controls. Representative inflammatory and myeloid-associated genes, including *Lyz2*, *Ly6c2* and *Ly6a2*, and the lymphoid-associated gene *Il7r* are highlighted. **M,** Trajectory inference of CD8⁺ T cell states in PBMCs from *Piezo1*^fl/fl^ and *Piezo1^Sost^* mice, showing redistribution of naive-like, effector and interferon-associated CD8⁺ T cell states. **N,** Chao1 richness index from paired TCR/BCR repertoire profiling of PBMCs from *Piezo1*^fl/fl^ and *Piezo1^Sost^* mice. **O,** D50 diversity index from paired TCR/BCR repertoire profiling of PBMCs from *Piezo1*^fl/fl^ and *Piezo1^Sost^*mice. For flow cytometry and serum cytokine analyses in **B–I**, n = 8 mice per group; each dot represents one mouse. For PBMC scRNA-seq analyses in **J–M**, n = 3 mice per group, with each mouse processed as an independent biological replicate. For paired TCR/BCR repertoire analyses in **N and O**, n = 3 mice per group, with each mouse processed as an independent biological replicate. Data are presented as mean ± s.e.m. *P* values are indicated in the plots. For comparisons among young, aged and aged+Load groups, one-way ANOVA followed by multiple-comparisons correction was used. For two-group comparisons, two-sided unpaired Student’s *t*-test was used.

To genetically validate the osteocyte mechanosensing axis, scRNA-seq of peripheral blood mononuclear cells (PBMCs) and profiling of the repertoire of T-cell receptors (TCRs) and B-cell receptors (BCRs) in *Piezo1^Sost^* mice were performed. The results of scRNA-seq of PBMCs resolved major immune lineages (Figure 5J), and compositional analysis revealed a shift toward an immune-aging-like profile with decreased lymphoid and increased myeloid fractions in *Piezo1^Sost^* mice (Figure 5K). Differential expression analysis of PBMC scRNA-seq data showed a myeloid/inflammatory shift in *Piezo1^Sost^* mice, with marked upregulation of *Lyz2*, *Ly6c2* and *Ly6a2* and reduced expression of the lymphoid-associated gene *Il7r* (Figure 5L). Within CD8^+^ T cells, naive-like and effector/interferon (IFN)-associated states were redistributed, and trajectory analysis further supported remodeling of CD8^+^ T cell state dynamics (Figure 5M). Repertoire profiling demonstrated impaired adaptive diversity (TCR/BCR) (Figure 5N and 5O). These findings underscore the mechano-bone-immune axis as a vital regulator of immune aging. Beyond facilitating acute defenses against infections, mechanical stimulation mediated by osteocyte *Piezo1* plays a crucial role in mitigating chronic inflammation, delaying systemic immunosenescence, and fostering immune homeostasis throughout life.

### PIEZO1 is evolutionarily conserved across species

Given the role of skeletal mechanosensing in bone and hematopoietic regulation, we next asked whether Piezo1, a major mechanosensitive ion channel, is conserved across species. Phylogenetic analysis showed that Piezo1 is broadly conserved among representative vertebrate lineages, including fish, amphibians, reptiles, birds, marsupials, placental mammals, and primates, with Drosophila included as an outgroup (Figure 6A). This conservation suggests that Piezo1-mediated mechanosensing is an evolutionarily maintained feature across diverse species.

**Figure 6.**
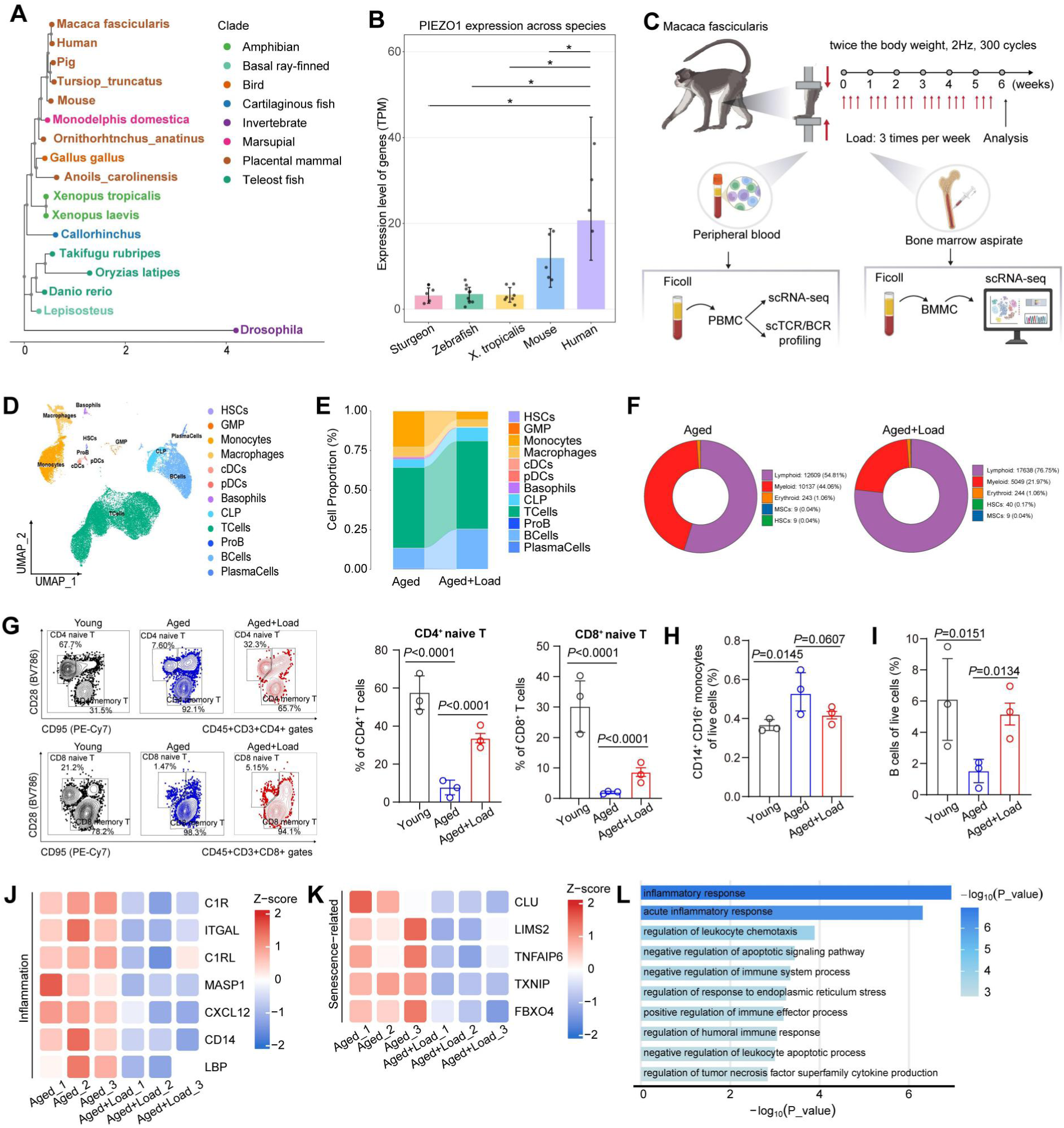
Evolutionary conservation of PIEZO1 and mechanical loading-induced immune remodeling in aged non-human primates. **A,** Maximum-likelihood phylogenetic analysis of PIEZO1 orthologous protein sequences across representative vertebrate species, with *Drosophila melanogaster* included as an invertebrate outgroup. Species are colored by major evolutionary clade. **B,** PIEZO1 expression across representative species based on publicly available RNA-seq datasets, shown as transcripts per million (TPM). **C,** Experimental scheme for dynamic long-bone loading in aged long-tailed macaques. Aged+Load macaques received mechanical loading equivalent to twice body weight at 2 Hz for 300 cycles per session, three sessions per week for 6 weeks. Age-matched sham-loaded macaques underwent the same experimental schedule without dynamic mechanical loading. Peripheral blood and bone marrow aspirates were collected for downstream immune profiling. **D,** UMAP visualization of major hematopoietic cell populations identified by scRNA-seq of bone marrow mononuclear cells from aged sham-loaded and aged+Load macaques. **E,** Relative proportions of major bone marrow hematopoietic populations in aged sham-loaded and aged+Load macaques based on scRNA-seq. **F,** Donut charts summarizing broad compartment-level distributions, including lymphoid, myeloid, erythroid, HSC and stromal-like compartments, in bone marrow from aged sham-loaded and aged+Load macaques (n = 3) . **G,** Representative flow cytometry plots and quantification of CD4⁺ naive T cells and CD8⁺ naive T cells in peripheral blood from young, aged sham-loaded and aged+Load macaques (n = 3). **H,** Quantification of CD14⁺CD16⁺ monocytes among live peripheral blood cells from young, aged sham-loaded and aged+Load macaques. **I,** Quantification of B cells among live peripheral blood cells from young, aged sham-loaded and aged+Load macaques. **J,** Heat map showing the relative abundance of inflammation-related proteins in bone marrow supernatants from aged sham-loaded and aged+Load macaques, as determined by proteomic analysis. **K,** Heat map showing the relative abundance of senescence-associated proteins in bone marrow supernatants from aged sham-loaded and aged+Load macaques, as determined by proteomic analysis. **L,** Functional enrichment analysis of proteins decreased in bone marrow supernatants after mechanical loading, highlighting inflammatory and immune-response-related pathways. For non-human primate experiments, young, aged sham-loaded and aged+Load macaques were analyzed as independent biological groups. Aged sham-loaded controls underwent the same anesthesia, positioning, limb fixation and session schedule as aged+Load macaques but did not receive dynamic mechanical loading. For bone marrow scRNA-seq and proteomic analyses, n = 3 biologically independent animals per group. For flow cytometry analyses in **G–I,** each dot represents one biologically independent animal. Data are presented as mean ± s.e.m. P values are indicated in the plots. For comparisons among young, aged sham-loaded and aged+Load groups, one-way ANOVA followed by multiple-comparisons correction was used unless otherwise indicated.

We then examined Piezo1 expression using publicly available transcriptomic datasets from representative species. Piezo1 was detectable across the species analyzed and showed higher expression in terrestrial mammals, including mouse and human, than in the aquatic vertebrates examined, including sturgeon and zebrafish, and in the amphibian Xenopus tropicalis (Figure 6B). Together with our functional studies in mice and non-human primates, these comparative analyses support the notion that skeletal mechanosensing is conserved across species and may contribute to the regulation of hematopoietic and immune homeostasis in mechanically active bone environments.

### Mechanical loading improves aged bone marrow hematopoiesis and peripheral immune imbalance in non-human primates

Building on our findings that skeletal mechanosensing is associated with immune regulation across species, we next explored whether mechanical loading could alleviate age-associated immune remodeling in higher mammals. To this end, we applied dynamic long-bone loading to aged long-tailed macaques (*Macaca fascicularis*) for 6 weeks (Figure 6C and Video 1). Loaded animals received mechanical stimulation equivalent to twice body weight at 2 Hz for 300 cycles per session, three sessions per week. Bone marrow aspirates and peripheral blood were collected from aged sham-loaded controls and aged loaded macaques, with young macaques included as reference controls for age-associated immune changes where indicated (Figure 6C). Bone marrow mononuclear cells were enriched and profiled by scRNA-seq (Figures 6C and S6A–S6B). This analysis resolved major hematopoietic populations in macaque bone marrow, including HSCs, GMPs, monocytes, macrophages, dendritic cells, lymphoid cells, B-lineage cells, and plasma cells (Figure 6D). Aged sham-loaded macaques showed a myeloid-skewed bone marrow composition, accompanied by reduced lymphoid representation. In aged loaded macaques, this imbalance was mitigated, with increased lymphoid representation and reduced myeloid representation relative to aged sham-loaded controls (Figures 6E and 6F). These findings suggest that mechanical loading restrains age-associated myeloid-biased hematopoietic remodeling in non-human primates.

Consistent with the bone marrow scRNA-seq data, peripheral blood flow cytometry showed that aging was associated with reduced CD28⁺CD95⁻ naive CD4⁺ and CD8⁺ T cell frequencies and reduced B cell frequencies, together with increased CD14⁺CD16⁺ monocytes. Compared with aged sham-loaded controls, aged loaded macaques showed increased naive CD4⁺ and CD8⁺ T cell frequencies and B cell frequencies, along with reduced CD14⁺CD16⁺ monocytes (Figures 6G–6I and S6C–S6D). Thus, mechanical loading partially corrected age-associated peripheral immune imbalance in aged macaques.

We next asked whether mechanical loading also altered the aged bone marrow microenvironment. Proteomic profiling of bone marrow supernatants revealed reduced abundance of multiple inflammation-related proteins in aged loaded macaques, including C1R, ITGAL, C1RL, MASP1, CXCL12, CD14, and LBP (Figure 6J). Several senescence-associated proteins, including CLU, LIMS2, TNFAIP6, TXNIP, and FBXO4, were also reduced after loading (Figure 6K). Enrichment analysis of proteins decreased by mechanical loading highlighted inflammatory and immune-response pathways, including inflammatory response, acute inflammatory response, regulation of leukocyte chemotaxis, and endoplasmic reticulum stress-associated pathways (Figure 6L). Together, these data indicate that mechanical loading remodels the aged macaque bone marrow niche toward a less inflammatory and less senescence-associated state, in parallel with improved hematopoietic and peripheral immune balance.

### Mechanical loading remodels adaptive immunity and restores immune repertoire integrity in aged non-human primates

To further determine whether bone-derived mechanical signals modulate peripheral adaptive immunity in primates, we profiled peripheral blood mononuclear cells from aged sham-loaded and aged loaded *Macaca fascicularis* by scRNA-seq. Integrated analysis resolved the major circulating immune lineages, including T cells, B cells, NK cells, monocytes, dendritic cells, neutrophils and platelets (Figure 7A). Compared with aged sham-loaded controls, aged loaded macaques showed an increased lymphoid fraction and a reduced myeloid fraction in peripheral blood (Figure 7B), consistent with the peripheral flow cytometry findings and the bone marrow remodeling observed after loading (Figure 6D–I). PBMC scRNA-seq further showed broad attenuation of inflammatory and myeloid-associated programs in aged loaded macaques, including reduced expression of S100A8 and S100A9, together with a shift toward a lymphoid-enriched and B-cell-associated composition and state (Figures S7A–S7G).

**Figure 7.**
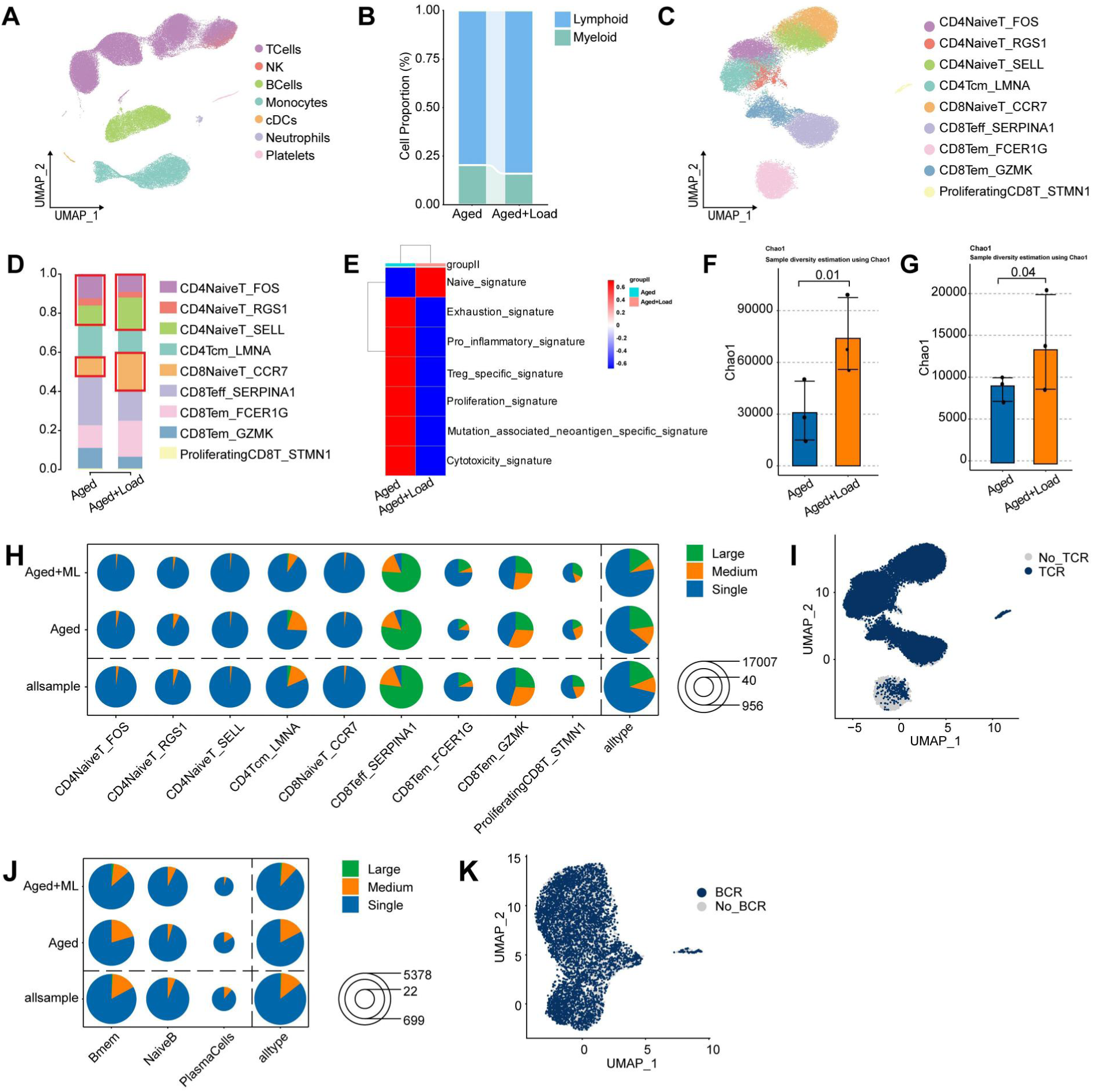
Mechanical loading remodels adaptive immune states and repertoire diversity in aged non-human primates. **A,** UMAP visualization of integrated peripheral blood mononuclear cell scRNA-seq profiles from aged sham-loaded and aged+Load macaques, colored by major circulating immune lineages. **B,** Relative proportions of lymphoid and myeloid compartments in aged sham-loaded and aged+Load macaques based on scRNA-seq. **C,** UMAP visualization of reclustered T cells from peripheral blood scRNA-seq, showing annotated naive-like, effector, interferon-associated and proliferative T cell states. **D,** Relative distribution of annotated T cell states among total T cells in aged sham-loaded and aged+Load macaques. **E,** Heatmap showing averaged gene-signature scores in T cells from aged sham-loaded and aged+Load macaques, including naive, exhaustion, pro-inflammatory, Treg, proliferation, mutation-associated neoantigen-specific and cytotoxicity signatures. **F,** Chao1 richness index of the TCR repertoire in aged sham-loaded and aged+Load macaques. **G,** Chao1 richness index of the BCR repertoire in aged sham-loaded and aged+Load macaques. **H,** Pie charts summarizing T cell clonotype size distributions across annotated T cell states in aged sham-loaded and aged+Load macaques. Colors denote clonotype expansion classes: single, medium and large. Pie size reflects the number of TCR-assigned cells. **I,** UMAP visualization highlighting cells with or without detected TCR sequences. **J,** Pie charts summarizing B cell clonotype size distributions across B cell subsets in aged sham-loaded and aged+Load macaques. Colors denote BCR clonotype expansion classes: single, medium and large. Pie size reflects the number of BCR-assigned cells. **K,** UMAP visualization highlighting cells with or without detected BCR sequences. Peripheral blood scRNA-seq and paired single-cell TCR/BCR repertoire profiling were performed using samples from aged sham-loaded and aged+Load macaques. Aged sham-loaded and aged+Load macaques were analyzed as independent biological groups. Unless otherwise indicated, *n* = 3 biologically independent macaques per group. Data are presented as mean ± s.e.m. *P* values are indicated in the plots. Comparisons between aged sham-loaded and aged+Load groups were performed using two-sided unpaired Student’s *t*-test unless otherwise indicated.

**Figure 8.**
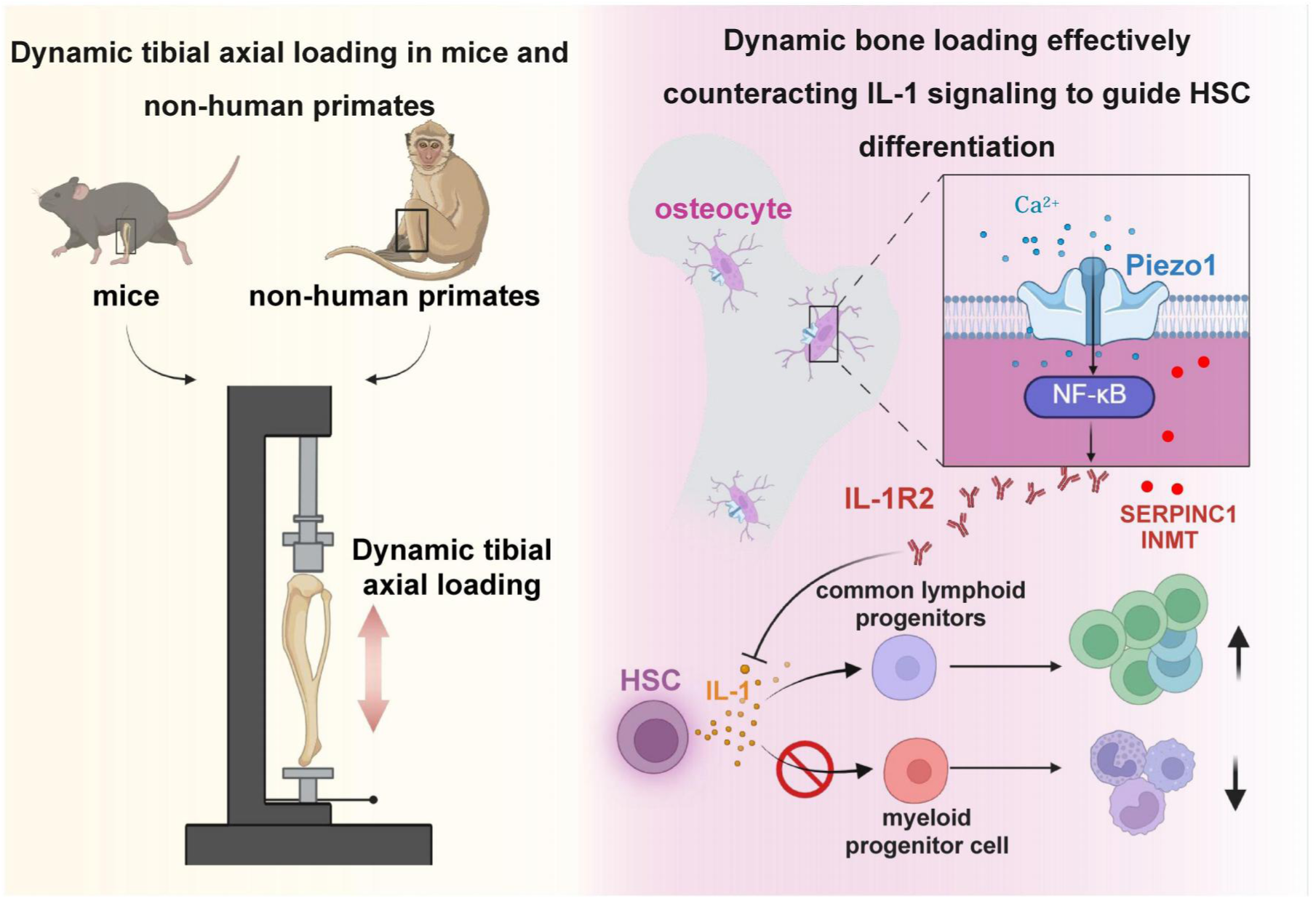
Dynamic bone loading activates an osteocyte Piezo1-dependent immunoregulatory molecular program to guide HSC fate and immune homeostasis. Schematic model illustrating the proposed mechano-bone-immune axis. Dynamic tibial axial loading was applied to mice and non-human primates to mimic physiological mechanical stimulation of bone. Mechanical loading is sensed by osteocytes through Piezo1-mediated mechanotransduction, which activates downstream signaling and induces a loading-responsive molecular program characterized by increased IL1R2, SERPINC1 and INMT. Among these factors, IL1R2 acts as a soluble decoy receptor to counteract IL-1 signaling and guide HSC differentiation, whereas SERPINC1 and INMT represent additional loading-responsive candidate molecules identified in the proteomic analysis. Together, these findings suggest that skeletal mechanical loading remodels the bone marrow microenvironment, restrains inflammatory cues and promotes HSC differentiation toward common lymphoid progenitors and lymphoid cell output while limiting excessive myeloid commitment. This model provides a conceptual framework by which skeletal mechanosensation contributes to immune homeostasis and limits immunosenescence. This figure was created with BioRender.com.

Changes in the adaptive immune receptor repertoire are a hallmark of immunosenescence, making TCR/BCR repertoire profiling a useful indicator of immune ageing and immune remodeling.^34,35^ We next reclustered T cells from the PBMC scRNA-seq dataset to examine how mechanical loading affects adaptive immune cell states. This analysis identified multiple naive-like, effector, interferon-associated and proliferative T-cell states (Figure 7C). Aged loaded macaques showed a shift toward increased naive-like T cell states and reduced effector or inflammatory T cell states relative to aged sham-loaded controls (Figure 7D). Consistently, gene-signature analysis showed increased naive-associated scores and reduced exhaustion-, pro-inflammatory-, proliferative- and cytotoxicity-associated scores in T cells from aged loaded macaques (Figure 7E). These findings suggest that mechanical loading remodels peripheral T-cell states toward a less inflammatory and more naive-like configuration in aged macaques.

We further performed paired single-cell TCR and BCR profiling to assess adaptive immune receptor repertoire diversity. Aged loaded macaques showed increased TCR and BCR repertoire richness, as reflected by higher Chao1 indices compared with aged sham-loaded controls (Figure 7F,G). Analysis of clonotype size distributions further showed that loading altered clonal architecture across T- and B-cell subsets, with changes in the relative proportions of single, medium and large clonotypes (Figure 7H,J). UMAP projection of cells with detectable TCR or BCR sequences confirmed broad coverage of T and B lineage compartments in the repertoire analysis (Figure 7I,K). Together, these data indicate that mechanical loading remodels adaptive immune states and improves immune receptor repertoire diversity in aged non-human primates.

Together with the bone marrow and peripheral immune changes described above, these findings support the concept that bone-derived mechanical signals can modulate adaptive immune architecture in primates and may provide a non-pharmacological approach for mitigating age-associated immune remodeling.

## Discussion

The findings of this study offer new insights into the complex relationship between osteocyte mechanosensing and the immune system, revealing an important, previously overlooked function of bone mechanosensing in shaping immune system dynamics. This discovery paves the way for potential therapeutic strategies aimed at preventing immunosenescence and addressing immune-related disorders. By adjusting the dynamic mechanical loading of bone, we could potentially develop innovative approaches to enhance immune health and resilience. Importantly, this study also indicates that the application of mechanical forces to bone tissue is not only essential for skeletal health but also significantly influences the evolutionary adaptation of the vertebrate immune system from aquatic to terrestrial environments, which may have been necessary to improve immune functionality in response to new challenges encountered in terrestrial habitats^36^.

The role of the skeletal system extends well beyond providing structural support and maintaining mineral homeostasis.^14^ Increasing evidence redefines bone as a vital immunological organ within the osteoimmune system,^37^ where continuous communication occurs between the skeletal and immune systems within the shared bone marrow microenvironment.^13^ This interaction is crucial for hematopoiesis and the development of immune cells,^38,39^ mediated by bone-derived cells that secrete niche factors (such as CXCL12 and SCF),^8,40^ cytokines (like IL-7),^28,41,42^ and other bone-derived factors^4^ that help regulate overall body homeostasis.^8,43,44^ However, the prevailing understanding has largely been limited to biochemical signaling. The inherent capacity as a mechanosensory tissue, as a fundamental aspect of bone biology, has been underappreciated in discussions of immune regulation. Our identification of the mechano-bone-immune axis introduces a crucial physical dimension to this relationship. We demonstrate that physiological mechanical forces, sensed by osteocytes through the Piezo1 channel, result in vital biochemical signaling via IL1R2 that directly influence the commitment of the hematopoietic stem cell lineage. This finding indicates that bone is not simply a passive scaffold or a source of soluble factors, but rather serves as an active mechanosensory interface that shapes systemic immunity in response to physical demands. Consequently, our work shifts the conceptual understanding of bone from merely facilitating energy and calcium metabolism and endocrine functions to a central coordinator that integrates mechanical stimuli for immune regulation, thereby offering a unified framework linking immune dysfunction, age-related immunosenescence, and immunity challenges associated with inactivity.

Complementing this emerging paradigm is the rapidly growing field of mechanoimmunology, which suggests that immune cells function as direct sensors of local tissue mechanics.^45^ Groundbreaking work has established that biophysical signals, such as extracellular matrix stiffness, fluid shear stress,^46^ and hydrostatic pressure,^47^ directly influence the effector functions of immune cells, including macrophages,^48^ dendritic cells,^49^ and lymphocytes.^50–52^ This modulation occurs through the activation of mechanosensitive pathways, including PIEZO channels,^53^ integrins,^54^ and YAP/TAZ,^46,48,50^ within the immune cells themselves. However, this conceptual framework has chiefly focused on how tissue-scale forces impact pre-existing, mature immune cells within localized environments during various conditions, such as infection,^53,55,56^ cancer,^51^ and fibrosis.^57^ A significant question remains: is there an organ level sensor that translates systemic and organism wide mechanical loads, such as gravitational force and exercise,^8^ into instructional cues that dictate the development and homeostatic set point of the immune system? Our discovery of the mechano-bone-immune axis provides an affirmative answer to this question and reveals a hierarchical mechanism for mechanical immunoregulation. While previous studies have primarily manipulated local fluid shear or substrate rigidity *in vitro* or targeted mechanosensors in specific immune lineages,^48,50,51,55^ our approach focuses on modulating comprehensive skeletal loading *in vivo*. We simulate conditions like DTAL, disuse (such as tail suspension), or selective deletion of mechanosensors in osteocytes. Our findings identified bone, specifically through its specialized mechanosensory osteocytes, as the primary orchestrator that translates these global mechanical cues into bioactive signals that direct hematopoietic fate decisions within the bone marrow. Thus, our work extends mechanoimmunology from a paradigm of local “tuning” of immune effector function to one of central programming of immune capacity by a dedicated mechanosensory organ. This establishes a crucial upstream layer of biomechanical control over immune processes, highlighting the importance of the skeletal system in immunity regulation at a systemic level.

The synchronous co-evolution of the skeletal and adaptive immune systems during the transition of vertebrates from aquatic to terrestrial habitats is postulated to reflect responses to multiple terrestrial pressures, including calcium scarcity,^58^ increased ultraviolet radiation,^18^ and a burgeoning array of pathogens.^59^ While the necessity for protection against ultraviolet radiation has been suggested as a reason for relocating the HSC niche into the BM, a concomitant and profound physical change (i.e., six-fold increase in gravitational force) has been largely overlooked. Our identification of the mechano-bone-immune axis positions the possibility of this ubiquitous biomechanical cue as an evolutionary driver: the gravitational loading may serve as a continuous, systemic signal that finely tuned hematopoietic output towards the lymphoid lineage, thus broadening the cellular foundation for adaptive immunity. This mechanism works alongside pivotal genetic innovations such as the RAG transposon, ^60,61^ which facilitated the generation of immune receptor diversity. Our findings highlight the strategic scaling of cellular capacity, ensuring robust lymphocyte production capable of deploying a diverse repertoire against terrestrial pathogens.^14,36^ Consequently, it appears that the mechanosensory function of osteocytes evolved not only for structural adaptation but also as a central regulator that links locomotive activity with the development of a sophisticated immune system suited for terrestrial life.

In summary, our study highlights a fundamental mechano-bone-immune axis, where mechanical forces are sensed by osteocytes and converted into biochemical signals that guide the fate of HSCs and program systemic immune homeostasis. This discovery defines bone as a key mechanosensory and immunoregulatory organ, revealing its vital role in linking physical activity to immune competence. The clinical implications of this axis are extensive and significant, offering a mechanistic basis for transforming therapeutic strategies, particularly to enhance immunity of individuals with restricted mobility, such as bedridden patients or astronauts experiencing microgravity. By demonstrating that mechanical loading on bone can directly combat immunosenescence and promote lymphoid-biased hematopoiesis, our findings advocate for a natural, safe, non-invasive, and non-pharmacological approach to bolster immune resilience against both acute and chronic infections, aging, and inflammation. Thus, our research provides a strong scientific foundation for the development of targeted devices that deliver controlled mechanical stimuli, paving the way for innovative therapies that warrant clinical trials for a range of immune-related diseases. Furthermore, the development of bone (osteocytes)-targeting Piezo1 agonists is also an important direction for the future.

## Resource availability

The sequencing datasets generated in this study, including mouse bone marrow scRNA-seq, mouse bone tissue bulk RNA-seq, mouse PBMC scRNA-seq with paired TCR/BCR repertoire sequencing, cynomolgus macaque PBMC scRNA-seq with paired TCR/BCR repertoire sequencing, and cynomolgus macaque bone marrow mononuclear cell scRNA-seq, have been deposited in the Genome Sequence Archive (GSA) of the National Genomics Data Center (NGDC) under accession numbers CRA039441, CRA043025, CRA043085, CRA043086, CRA043087, CRA043129, CRA043130, CRA043131, and CRA043132. The mass spectrometry proteomics data have been deposited to the ProteomeXchange Consortium via the iProX partner repository under accession number PXD048706. The cynomolgus macaque bone marrow supernatant proteomics dataset has been deposited in the OMIX database of the National Genomics Data Center (NGDC) under accession number OMIX017116.

This paper does not report original code. Additional information required to reanalyze the data reported in this paper is available from the lead contact upon reasonable request.

## Acknowledgements

This work was supported by the National Natural Science Foundation of China (Nos. 82530080, 92268204, 32300965, and 82522003).

## Author contributions

X.B., X.Y. and Y.L. conceived the project, designed and interpreted the experiments. W.Z., Y.P., X.X., and J.D. performed animal and cell experiments. H.Z., Z.Y., S.Z., and X.L. generated and characterized the genetically modified mice. X.Y., J.H., H.J., and D.C. performed single-cell RNA sequencing. W.Z., Y.P., and X.X. performed bioinformatic and statistical analyses. X.B. and X.Y. wrote the manuscript.

## Methods

### Mice experiments

Male C57BL/6J mice were purchased from Guangdong Weitong Lihua Laboratory Animal Technology Co., Ltd. (Foshan, China). Unless otherwise stated, male mice were used for all mouse experiments. Young wild-type mice were used at 12 weeks of age, and aged wild-type mice were used at 21 months of age. *Sost*-CreERT2 mice were obtained from Shanghai Model Organisms Center, Inc. (Shanghai, China; Cat# NM-KI-200171). *Dmp1*-Cre mice were obtained from The Jackson Laboratory (B6N. FVB-Tg(*Dmp1*-cre)1Jqfe/BwdJ; Stock No. 023047; RRID: IMSR_JAX:023047). *Il1r2*^fl/fl^ mice were obtained from GemPharmatech (Strain No. T051901). *Piezo1*^fl/fl^ mice were obtained from the UC Davis KOMP Repository. All mouse lines were maintained on a C57BL/6J background.

Osteocyte-lineage *Piezo1*-deficient mice were generated by crossing *Piezo1*^fl/fl^ mice with *Dmp1*-Cre mice to obtain *Dmp1*-Cre ; *Piezo1*^fl/fl^ mice, hereafter referred to as *Piezo1^Dmp1^* mice. Cre-negative *Piezo1*^fl/fl^ littermates were used as controls. Inducible osteocyte-lineage Piezo1-deficient mice were generated by crossing *Piezo1*^fl/fl^ mice with *Sost*-CreERT2 mice to obtain *Sost*-CreERT2; *Piezo1*^fl/fl^ mice, hereafter referred to as *Piezo1^Sost^* mice. Cre-negative *Piezo1*^fl/fl^ littermates were used as controls. Inducible osteocyte-lineage *Il1r2*-deficient mice were generated by crossing *Il1r2*^fl/fl^ mice with *Sost*-CreERT2 mice to obtain *Sost*-CreERT2; *Il1r2*^fl/fl^ mice, hereafter referred to as *Il1r2^Sost^* mice. Cre-negative *Il1r2*^fl/fl^ littermates were used as controls. All conditional knockout mice and littermate controls used in this study were male.

For *Sost*-CreERT2-mediated inducible deletion in osteocyte-lineage cells, 3-month-old *Piezo1^Sost^* or *Il1r2^Sost^*mice were administered tamoxifen. Tamoxifen was dissolved in corn oil at 20 mg/ml and administered by intraperitoneal injection at 0.2 mg/g body weight per dose once daily for 5 consecutive days. Cre-negative *Piezo1*^fl/fl^ or *Il1r2*^fl/fl^ littermate controls received the same tamoxifen regimen. Experiments were initiated 7 days after the final tamoxifen administration unless otherwise indicated. Deletion efficiency of *Piezo1* or *Il1r2* in osteocyte-lineage cells was verified by immunofluorescence staining and quantitative PCR analysis of isolated osteocyte-enriched bone fractions, as described below. Genotyping was performed by PCR using genomic DNA isolated from tail or ear biopsies. Primer sequences are listed in the Key Resources Table.

Mice were housed under specific pathogen-free conditions in plastic cages at 22 ± 1 °C under a 12-h light/dark cycle and were provided standard rodent chow and water ad libitum throughout the study. Mice were randomly assigned to experimental groups where applicable. Investigators were blinded to group allocation during histological scoring, flow cytometric gating and image quantification where applicable. All mouse experiments were performed in accordance with institutional guidelines and were approved by the Animal Ethics Committee of Southern Medical University under approval number SMUL2020141. For sepsis experiments, including cecal ligation and puncture and LPS-induced systemic inflammation models, mice were monitored closely after challenge, with increased monitoring frequency during the acute phase. Humane endpoint criteria were predefined before the experiments. Mice were euthanized if they showed moribund status, loss of righting reflex, inability to ambulate or access food and water, persistent lateral recumbency, severe respiratory distress, severe hypothermia, unresponsiveness to external stimuli, or other signs of severe and irreversible distress. Animals that reached humane endpoints were euthanized immediately using an institutionally approved method and were counted as deaths at the time of euthanasia in Kaplan–Meier survival analyses.

### Aged cynomolgus macaque mechanical loading

Young adult male long-tailed macaques (*Macaca fascicularis*; 8 years old; n = 3) were included as youthful reference controls for peripheral blood flow-cytometry analyses. Aged male long-tailed macaques (*Macaca fascicularis*; 21 years old) were assigned to an aged sham-loaded control group (n = 3) or an aged mechanical-loading group (aged+Load; n = 3). Aged sham-loaded macaques underwent the same anesthesia, positioning, limb fixation and session schedule without dynamic mechanical loading, whereas aged+Load macaques were subjected to dynamic long-bone loading for 6 weeks. Thus, aged sham-loaded and aged+Load macaques were analyzed as independent parallel groups.

For each loading session, animals were anesthetized with isoflurane. Anesthesia was induced with 3%–4% isoflurane and maintained at approximately 3% isoflurane during the procedure. The tibiae were secured in custom clamps coupled to an electrodynamic loading system (ElectroForce 3510; TA Instruments). In the aged+Load group, each tibia was loaded at 2 Hz with a peak compressive load equivalent to twice body weight for 300 cycles per limb per session, three sessions per week for 6 weeks. In the aged sham-loaded group, animals underwent the same anesthesia, positioning, limb fixation and session schedule, but no dynamic load was applied.

Physiological parameters, including ECG, SpO₂, end-tidal CO₂ and body temperature, were continuously monitored throughout the procedure. During anesthesia recovery, animals were kept warm and monitored until full recovery from anesthesia. Animals were further monitored throughout the intervention period by trained personnel under veterinary supervision. No local injury, lameness, body-weight loss or abnormal behavior was observed after mechanical loading. Loading was not performed, or was stopped, if animals showed signs of unstable anesthesia, abnormal respiration, impaired recovery, limb injury, persistent pain, swelling or other procedure-related adverse effects.

Bone marrow aspirates were collected from the tibial plateau under anesthesia and aseptic conditions from aged sham-loaded and aged+Load macaques for bone marrow mononuclear cell scRNA-seq and proteomic analysis of bone marrow supernatants. Peripheral blood samples from aged sham-loaded and aged+Load macaques were used for PBMC scRNA-seq and paired single-cell TCR/BCR repertoire profiling. Peripheral blood samples from young control, aged sham-loaded and aged+Load macaques were used for flow-cytometry analyses, as indicated. All non-human primate procedures were approved by the Animal Experimental Ethics Committee of the Institute of Zoology, Guangdong Academy of Sciences under approval number GIZ20241125 and were performed in accordance with institutional guidelines for the care and use of laboratory animals.

### DTAL in mice

Mechanical loading was performed bilaterally using an electrodynamic test instrument [ElectroForce 3300; TA Instruments, Eden Prairie, MN, USA] with custom tibial clamps as previously described for murine tibial loading^62^. Mice were anesthetized with 1.5–2.0% isoflurane and positioned with the knee and ankle aligned along the loading axis. A small pre-load of 0.2 N was applied to ensure stable contact. Mechanical loading was delivered using a sinusoidal waveform at 2 Hz, with a peak compressive load of 2 N and 300 cycles per limb per session. Unless otherwise stated, loading was performed three times per week for 2 weeks. For aged mouse experiments, loading was performed three times per week for 4 weeks. Sham-loaded mice underwent identical anesthesia, positioning and limb fixation without dynamic load application. Respiration and reflex responses were monitored throughout anesthesia. Because the procedure was non-invasive and performed under brief anesthesia, analgesics were not routinely administered. Mice were monitored after each session for signs of pain, impaired ambulation, limb swelling or injury.

### Flow cytometry

Mice were euthanized according to institutional guidelines. Bone marrow cells were isolated from femurs and tibiae by flushing the marrow cavity with ice-cold FACS buffer consisting of PBS supplemented with 2% fetal bovine serum (FBS). Cell suspensions were mechanically dissociated, filtered through a 70-μm cell strainer and centrifuged at 400 × g for 5 min at 4°C. Red blood cells were lysed with ammonium-chloride-potassium (ACK) lysing buffer (Thermo Fisher Scientific, Cat# A1049201) for 10 min at room temperature. Cells were washed twice with FACS buffer and passed through a 40-μm cell strainer before staining.

Peripheral blood was collected by cardiac puncture using heparin-coated syringes and transferred into EDTA-containing tubes. Spleens were dissected aseptically and mechanically dissociated through 70-μm cell strainers. Red blood cells in peripheral blood and splenic cell suspensions were lysed with ACK lysing buffer. Cells were washed twice with cold FACS buffer and filtered through 40-μm cell strainers. Cell number and viability were assessed by trypan blue exclusion before antibody staining.

For surface staining, single-cell suspensions were incubated with viability dye to exclude dead cells and then stained with fluorophore-conjugated antibodies for 30 min at 4°C in the dark. Fc receptor blocking with anti-CD16/CD32 antibody was performed for staining panels that did not use CD16/32 as an analytical marker. For hematopoietic stem and progenitor cell panels in which CD16/32 was used to define GMPs, Fc receptor blocking with anti-CD16/CD32 was omitted to avoid interference with FcγR staining. Antibodies used for flow cytometry are listed in the Key Resources Table.

For bone marrow hematopoietic stem and progenitor cell analysis, debris, doublets and dead cells were excluded before gating. Lineage-negative cells were identified using a lineage antibody cocktail. LSK cells were defined as Lin⁻Sca-1⁺c-Kit⁺ cells. CLPs were gated as Lin⁻Sca-1^lo^c-Kit^lo^IL-7Rα⁺Flt3⁺ cells. GMPs were gated within the LK compartment as Lin⁻Sca-1⁻c-Kit⁺CD34⁺FcγR^hi^ cells.

For peripheral blood and spleen immune profiling, CD45⁺ leukocytes were analyzed after exclusion of debris, doublets and dead cells. Myeloid cells were identified as CD11b⁺ cells, neutrophils as CD11b⁺Ly6G⁺ cells and Ly6C^hi^ monocytes as CD11b⁺Ly6C^hi^ cells. B cells were identified as CD19⁺B220⁺ cells in general immune profiling. For age-associated B cell analysis, B-lineage cells were gated as CD19⁺IgM⁺ cells, mature B cells as CD43⁻CD93⁻ cells, and age-associated B cells were defined based on CD23 and CD21/CD35 expression as indicated in the gating strategy. T cells were identified as CD3⁺ cells. Naive, central memory-like and effector memory-like T cell populations were defined based on CD44 and CD62L expression, with naive T cells gated as CD44^lo^CD62L^hi^ cells.

Flow cytometry was performed using a CytoFLEX flow cytometer or BD LSRFortessa X-20, and cell sorting was performed using a MoFlo XDP cell sorter where indicated. Data were analyzed using FlowJo v10. Gates were set using fluorescence-minus-one controls where appropriate, and the same gating strategy was applied across experimental groups within each experiment.

### Induction of Polymicrobial Sepsis

Polymicrobial sepsis was induced via cecal ligation and puncture according to previously described protocols with minor modifications. Mice were anesthetized with isoflurane and placed on a heating pad to maintain homeostatic body temperature. Following abdominal depilation, the surgical site was disinfected using povidone-iodine followed by 70% ethanol. Under aseptic conditions, a midline laparotomy was performed to exteriorize the cecum.

To achieve mid-grade sepsis, the cecum was ligated at approximately 50% of its length from the distal end using 4-0 silk suture, ensuring that intestinal continuity was maintained to avoid bowel obstruction. The ligated segment was then subjected to a single through-and-through puncture using an 18-gauge sterile needle. A small amount of extralumenal fecal matter was gently extruded to confirm patency of the punctures. The cecum was then repositioned into the abdominal cavity. The incision was closed in two layers: the peritoneum and musculature were secured with 5-0 absorbable continuous sutures, and the skin was closed with 5-0 silk interrupted sutures.

### Reverse transcription and quantitative PCR

Total RNA was extracted from isolated cells or tissue samples using TRIzol reagent or the indicated RNA extraction kit according to the manufacturer’s instructions. RNA concentration and purity were assessed using a NanoDrop spectrophotometer. Equal amounts of total RNA were reverse-transcribed into complementary DNA (cDNA) using the High-Capacity cDNA Reverse Transcription Kit (Thermo Fisher Scientific) according to the manufacturer’s instructions.

Quantitative PCR was performed using SYBR Fast qPCR Master Mix (Kapa Biosystems) on a CFX384 Real-Time PCR Detection System (Bio-Rad). Each reaction was performed in technical triplicate. Relative gene expression was normalized to Gapdh and calculated using the 2^−ΔΔCt^ method. Primer sequences are listed in the Key Resources Table.

### Tissue Processing and Immunofluorescence

Tissues were fixed in 4% paraformaldehyde (PFA) at 4°C for 24 h, followed by paraffin embedding and sectioning at a thickness of 3 µm. Following deparaffinization in xylene and rehydration through a graded ethanol series, heat-induced epitope retrieval (HIER) was performed by incubating sections in 10 mM sodium citrate buffer (pH 6.0) overnight at 60°C. To minimize non-specific binding, sections were blocked with 1% normal goat serum in PBS for 1 h at room temperature.

Sections were incubated overnight at 4°C with the following primary antibodies: goat anti-SOST/Sclerostin (R&D Systems, AF1589; 1:50), rabbit anti-PIEZO1 (HUABIO, HA601100; 1:100), mouse anti-IL-1R2 (Proteintech, 60262-1-Ig; 1:100), and goat anti-DMP-1 (R&D Systems, AF4386; 1:100). After three 5-min washes in PBS, sections were incubated for 1 h at 37°C with species-specific Alexa Fluor-conjugated secondary antibodies (Invitrogen/Thermo Fisher Scientific), including donkey anti-rabbit 488 (A-21206; 1:500), donkey anti-mouse 488 (A-21202; 1:1,000), donkey anti-mouse 594 (A-21203; 1:500), donkey anti-goat 488 (A-11055; 1:500), and donkey anti-sheep 594 (A-11016; 1:500). Nuclei were counterstained with DAPI (1:500). Sections were mounted in 90% glycerol and stored at 4°C protected from light.

Immunofluorescence images were acquired using an Olympus FV3000 confocal laser scanning microscope. Laser power and gain settings were maintained constant across all samples within an experiment to ensure comparability.

### Serum Cytokine Quantification by ELISA

Blood was collected via cardiac puncture 6 h post-induction of sepsis. To obtain serum, whole blood was allowed to clot at room temperature for 2 h, followed by centrifugation at 400 × g for 10 min at 4°C. The supernatant was aliquoted and stored at –80°C until further analysis. Serum concentrations of IL-6, IL-1β, and TNF-α were determined using highly sensitive, precoated ELISA kits (Dakewe Biotech, Shenzhen, China; IDs: #1210602, #1210122, and #1217202, respectively) according to the manufacturer’s instructions.

Briefly, serum samples, standards, and blank controls were incubated with biotinylated detection antibodies at 37°C for 90 min. Following four washes with wash buffer to remove unbound proteins, Streptavidin-HRP working solution was added and incubated at 37°C for 30 min. After a second wash cycle, 3,3’,5,5’-tetramethylbenzidine (TMB) substrate was added and incubated in the dark at 37°C for 10 min for chromogenic development. The reaction was terminated using an acidic Stop Solution. The optical density (OD) was measured at 450 nm (with wavelength correction set to 570 nm or 630 nm) using a multimodal microplate reader. Cytokine concentrations were calculated by interpolating the OD values against a four-parameter logistic (4-PL) standard curve. All samples were processed in technical duplicates.

## Protein extraction and immunoblotting

Tissues or cell pellets were lysed in ice-cold RIPA buffer supplemented with 1× protease and phosphatase inhibitor cocktail (Thermo Fisher Scientific). Lysates were mechanically homogenized, sonicated and centrifuged at 12,000 × g for 15 min at 4°C to remove insoluble debris. Protein concentrations were determined using a BCA Protein Assay Kit (Pierce) according to the manufacturer’s instructions. Equal amounts of protein were mixed with Laemmli sample buffer, denatured at 100°C for 5 min and separated by SDS-PAGE. Proteins were transferred onto 0.22-μm nitrocellulose membranes (Millipore) at 300 mA for 90 min at 4°C. Membranes were blocked for 1 h at room temperature in 5% non-fat milk in TBST for total protein detection or 5% BSA in TBST for phosphorylated protein detection. Membranes were then incubated overnight at 4°C with primary antibodies against IL1R2 (Proteintech, Cat# 60262-1-Ig; 1:1,000), NF-κB p65 (Cell Signaling Technology, Cat# 8242; 1:1,000), phospho-NF-κB p65 Ser536 (Cell Signaling Technology, Cat# 3033; 1:1,000) and GAPDH (Cell Signaling Technology, Cat# 97166; 1:5,000). After three washes in TBST, membranes were incubated with HRP-conjugated secondary antibodies for 1 h at room temperature. Protein bands were detected using enhanced chemiluminescence substrate (Bio-Rad) and imaged with a ChemiDoc Imaging System (Bio-Rad). Band intensities were quantified using ImageJ. IL1R2 and total NF-κB p65 were normalized to GAPDH, whereas phospho-NF-κB p65 was normalized to total NF-κB p65. Representative immunoblots from at least three independent experiments are shown unless otherwise indicated.

### Lung histopathology and injury scoring

To assess sepsis-induced lung injury, left lung lobes were fixed in 4% paraformaldehyde, embedded in paraffin and sectioned at 3 μm thickness for H&E staining. Histopathological changes were evaluated by two independent investigators blinded to experimental group allocation. Lung injury was scored using a semi-quantitative system based on four parameters: alveolar congestion, hemorrhage, neutrophil infiltration or aggregation in the airspace or vessel wall, and alveolar wall thickening or hyaline membrane formation. Each parameter was graded from 0 to 4 according to the severity and extent of injury: 0, no injury or minimal damage; 1, mild injury affecting <25% of the field; 2, moderate injury affecting 25%–50% of the field; 3, severe injury affecting 50%–75% of the field; and 4, extremely severe injury affecting >75% of the field. The total lung injury score was calculated as the sum of the four individual parameter scores, with a maximum score of 16. For each lung section, five randomly selected non-overlapping fields were examined at 200× magnification. The average score from the five fields was calculated for each mouse and used as one biological replicate for statistical analysis.

### Systemic administration of recombinant IL1R2

To evaluate the in vivo effect of IL1R2 on the bone marrow compartment, mice were administered recombinant mouse IL1R2 (MCE, Cat# HY-P72568). Lyophilized recombinant IL1R2 was reconstituted in sterile PBS containing 0.1% endotoxin-free bovine serum albumin (BSA). Mice received recombinant IL1R2 at 5 mg/kg body weight, corresponding to approximately 100 μg per 20-g mouse, by tail-vein injection in a total volume of 100 μl. Vehicle control mice received an equal volume of sterile PBS containing 0.1% endotoxin-free BSA.

For longitudinal administration, recombinant IL1R2 or vehicle was administered by tail-vein injection three times per week for 2 consecutive weeks. Femurs and tibiae were harvested 24 h after the final injection for bone marrow analysis.

### PBMC isolation

Peripheral blood was collected into EDTA-containing tubes and processed immediately. Peripheral blood mononuclear cells (PBMCs) were isolated by density-gradient centrifugation using Ficoll-based separation medium according to the manufacturer’s instructions. The mononuclear cell layer was collected, washed with PBS, and treated with red blood cell lysis buffer when residual erythrocytes were present. Cells were then washed twice with PBS and filtered through a 40-μm cell strainer to obtain a single-cell suspension.

Cell concentration and viability were determined by trypan blue exclusion. For single-cell RNA-seq and paired single-cell TCR/BCR library preparation, cells were resuspended in PBS containing 0.04% BSA and adjusted to the concentration required by the single-cell processing platform. Only samples with cell viability greater than 90% were used for single-cell library preparation. Samples with excessive cell debris or aggregation were excluded or reprocessed according to predefined quality-control criteria before library preparation.

### Single-cell RNA sequencing and immune repertoire sequencing

Single-cell RNA sequencing was performed on mouse bone marrow and cynomolgus macaque bone marrow samples using the 10x Genomics Chromium platform. Single-cell RNA sequencing with paired immune repertoire sequencing was performed on mouse peripheral blood mononuclear cells and cynomolgus macaque peripheral blood mononuclear cells using the GEXSCOPE Single Cell Immuno-TCR/BCR Kit (Singleron Biotechnologies). Libraries were sequenced on an Illumina platform using paired-end sequencing.

For bone marrow samples, single-cell suspensions were loaded onto the 10x Genomics Chromium Controller according to the manufacturer’s instructions to generate gel bead-in-emulsions. Cell barcodes and unique molecular identifiers were introduced during reverse transcription, enabling transcriptomic reads to be assigned to individual cells. Barcoded cDNA was recovered, amplified and used for gene expression library construction according to the 10x Genomics protocol.

For peripheral blood samples, single-cell suspensions were processed using the GEXSCOPE Single Cell Immuno-TCR/BCR workflow according to the manufacturer’s instructions. Briefly, cells were loaded onto microfluidic chips for single-cell capture and barcoding. After cell lysis, mRNA and immune receptor transcripts were captured and reverse-transcribed to generate barcoded cDNA. Gene expression libraries were constructed from amplified cDNA. In parallel, TCR and BCR transcripts were enriched from the remaining cDNA by targeted amplification, followed by PCR amplification and library construction to generate paired single-cell TCR-seq and BCR-seq libraries.

Final libraries were subjected to quality control and sequenced on an Illumina platform. Library construction, quality assessment and sequencing were performed according to the manufacturers’ protocols.

### Processing and analysis of single-cell RNA-seq and immune repertoire data

Raw sequencing data were processed using platform-specific pipelines. For 10x Genomics bone marrow scRNA-seq libraries, raw base-call files were demultiplexed and converted to FASTQ files, and gene-cell count matrices were generated by aligning reads to the corresponding reference genomes. Mouse reads were aligned to the GRCm39 reference genome, and cynomolgus macaque reads were aligned to the Macaca_fascicularis_6.0 reference genome. Cell barcodes and unique molecular identifiers (UMIs) were used to assign reads to individual cells and collapse PCR duplicates.

For Singleron GEXSCOPE peripheral blood scRNA-seq libraries, raw sequencing data were processed using CeleScope v2.7.3 (Singleron Biotechnologies). Cell barcodes and UMIs were extracted and corrected from R1 reads, and adaptor sequences and poly(A) tails were trimmed from R2 reads. Clean reads were aligned to the corresponding reference genomes using STARSolo in STAR v2.7.11a. Reads with the same cell barcode, UMI and gene annotation were collapsed to generate gene-cell count matrices.

Single-cell gene expression matrices were analyzed using Scanpy v1.9.3 in Python 3.10 and CeleLens Cloud (Singleron Biotechnologies), as appropriate. Low-quality cells and low-abundance genes were removed based on the distributions of detected genes, UMI counts and mitochondrial transcript fractions for each dataset. After library-size normalization and logarithmic transformation, highly variable genes were identified using the Seurat v3 method. Principal component analysis was performed using highly variable genes, followed by neighborhood graph construction, Louvain clustering and visualization by uniform manifold approximation and projection (UMAP). Cell types were annotated based on canonical marker genes, automated annotation results and manual curation.

Differentially expressed genes between experimental groups or cell clusters were identified using the built-in functions of Scanpy or CeleLens Cloud. Functional enrichment analysis was performed for selected differentially expressed genes as indicated in the figure legends. Additional downstream analyses, including cell-state scoring and immune-lineage composition analysis, were performed using the corresponding modules in Scanpy or CeleLens Cloud.

For immune repertoire analysis of peripheral blood samples, paired single-cell TCR and BCR data were processed using the Singleron immune repertoire analysis workflow. Productive TCR and BCR contigs were assigned to individual cell barcodes and integrated with the corresponding single-cell gene expression data. For TCR analysis, cells with productive TRA and TRB chains were retained, and each unique TRA–TRB pair was defined as a clonotype. For BCR analysis, cells with productive IGH and IGK or IGL chains were retained, and each unique IGH–IGK/IGL pair was defined as a clonotype. Clonotypes detected in at least two cells were considered clonally expanded. Repertoire richness and clonotype-size distributions were analyzed as indicated in the figure legends.

### Phylogenetic analysis of PIEZO1

PIEZO1 orthologous protein sequences were retrieved from NCBI for representative species spanning major metazoan and vertebrate clades, including the invertebrate outgroup Drosophila melanogaster; cartilaginous fish, Callorhinchus milii; basal ray-finned fish, Lepisosteus oculatus; teleost fish, Danio rerio, Oryzias latipes and Takifugu rubripes; amphibians, Xenopus tropicalis and Xenopus laevis; reptiles, Anolis carolinensis; birds, Gallus gallus; monotremes, Ornithorhynchus anatinus; marsupials, Monodelphis domestica; and placental mammals, Homo sapiens, Macaca fascicularis, Mus musculus, Sus scrofa and Tursiops truncatus. Protein sequences were aligned using MUSCLE v5.1 with default parameters. Poorly aligned regions were trimmed using trimAl v1.4 with the automated1 strategy. A maximum-likelihood phylogenetic tree was reconstructed using IQ-TREE v2.2.0 under the best-fit model selected by ModelFinder. Branch support was assessed using 1,000 ultrafast bootstrap replicates. The tree was rooted using Drosophila melanogaster as the outgroup and visualized using ggtree.

### Gene expression analysis of PIEZO1

Publicly available RNA-seq datasets were obtained from the Gene Expression Omnibus for representative skeletal or skeletal-related tissues or cell populations from sturgeon, zebrafish, Xenopus tropicalis, mouse and human. The datasets included sturgeon scale samples (GSE235280), zebrafish skeletal cell populations (GSE267333), Xenopus tropicalis craniofacial skeletal cell populations (GSE231325), mouse metatarsal bone samples (GSE284187) and human phalanx samples (GSE286924). For each dataset, transcript abundance was calculated as transcripts per million (TPM) from raw counts using gene length information, followed by log2(TPM + 1) transformation. PIEZO1 orthologs were identified using a precompiled ortholog table.

Differences in log2(TPM + 1) values among species were assessed using Welch’s ANOVA followed by Games–Howell post hoc tests. P values were adjusted for multiple comparisons using the Benjamini–Hochberg method, and adjusted P < 0.05 was considered statistically significant. Because these RNA-seq datasets were generated independently across different species and experimental platforms, cross-species expression comparisons were interpreted as comparative transcriptomic analyses rather than direct quantitative measurements across species.

### Statistics and reproducibility

No statistical method was used to predetermine sample size. Sample sizes were chosen based on previous studies using similar animal models and were sufficient to detect robust biological effects in the primary endpoints. Animals were randomly assigned to experimental groups where applicable. For genetic mouse models, littermate controls were used whenever possible. Investigators were blinded to group allocation during histological scoring, flow-cytometry gating and image quantification where applicable. No data points were excluded from analysis unless predefined exclusion criteria described in the relevant Methods sections were met.

Statistical analyses were performed using GraphPad Prism 8.0 unless otherwise indicated. Data are presented as mean ± s.e.m. unless stated otherwise. Each dot in the plots represents one biologically independent animal unless otherwise indicated. For two-group comparisons, two-sided unpaired Student’s t-test was used when data were approximately normally distributed. For comparisons involving more than two groups, one-way ANOVA followed by multiple-comparisons correction was used. For experiments involving two independent variables, such as genotype and mechanical loading, two-way ANOVA followed by multiple-comparisons correction was used. For ordinal histopathological scores, Mann–Whitney U test or Kruskal–Wallis test with multiple-comparisons correction was used as appropriate. Survival curves were analyzed using the log-rank Mantel–Cox test.

For non-human primate experiments, young control, aged sham-loaded and aged+Load macaques were analyzed as independent biological groups. Young versus aged sham-loaded comparisons were used to assess age-associated changes where young macaques were included, whereas aged sham-loaded versus aged+Load comparisons were used to assess the effect of mechanical loading. Aged sham-loaded animals underwent the same anesthesia, positioning, limb fixation and experimental schedule without dynamic mechanical loading. Young macaques were included as reference controls for peripheral blood flow-cytometry analyses where indicated. Bone marrow scRNA-seq, peripheral blood scRNA-seq, paired scTCR/scBCR repertoire profiling and bone marrow supernatant proteomic analyses in macaques were performed using aged sham-loaded and aged+Load animals.

For single-cell RNA-seq and paired TCR/BCR repertoire analyses, the biological replicate was defined as an individual mouse or macaque, not an individual cell or clonotype. Mouse bone marrow scRNA-seq analyses in Figures 1 and 2 were performed using *n* = 2 mice per group, with each mouse processed independently for library preparation and sequencing. Mouse PBMC scRNA-seq and paired TCR/BCR repertoire analyses were performed using *n* = 3 mice per group, with each mouse processed as an independent biological replicate. Non-human primate bone marrow mononuclear cell scRNA-seq, PBMC scRNA-seq and paired scTCR/scBCR repertoire analyses were performed using *n* = 3 biologically independent aged sham-loaded and *n* = 3 biologically independent aged+Load macaques unless otherwise indicated. Cell-level analyses, including clustering, marker-gene visualization, cell-state scoring and cell-composition analysis, were used to characterize cellular states and were interpreted together with animal-level comparisons and validation experiments.

For bulk RNA-seq, proteomic and differential gene-expression analyses, normalization procedures, statistical methods and multiple-testing correction are described in the relevant Methods sections. Differential features were considered significant using the indicated nominal *P* value or false discovery rate threshold, as specified for each analysis. Exact *P* values, sample sizes and statistical tests are provided in the figure legends or Methods. *P* values < 0.05 were considered statistically significant.

