## Supplementary Materials for "Bone Mechanosensing Dictates Hematopoietic Stem Cell Fate and Immune Homeostasis"

The PDF file includes:

Figure S1 to Figure S7

Supplemental Figure Legends

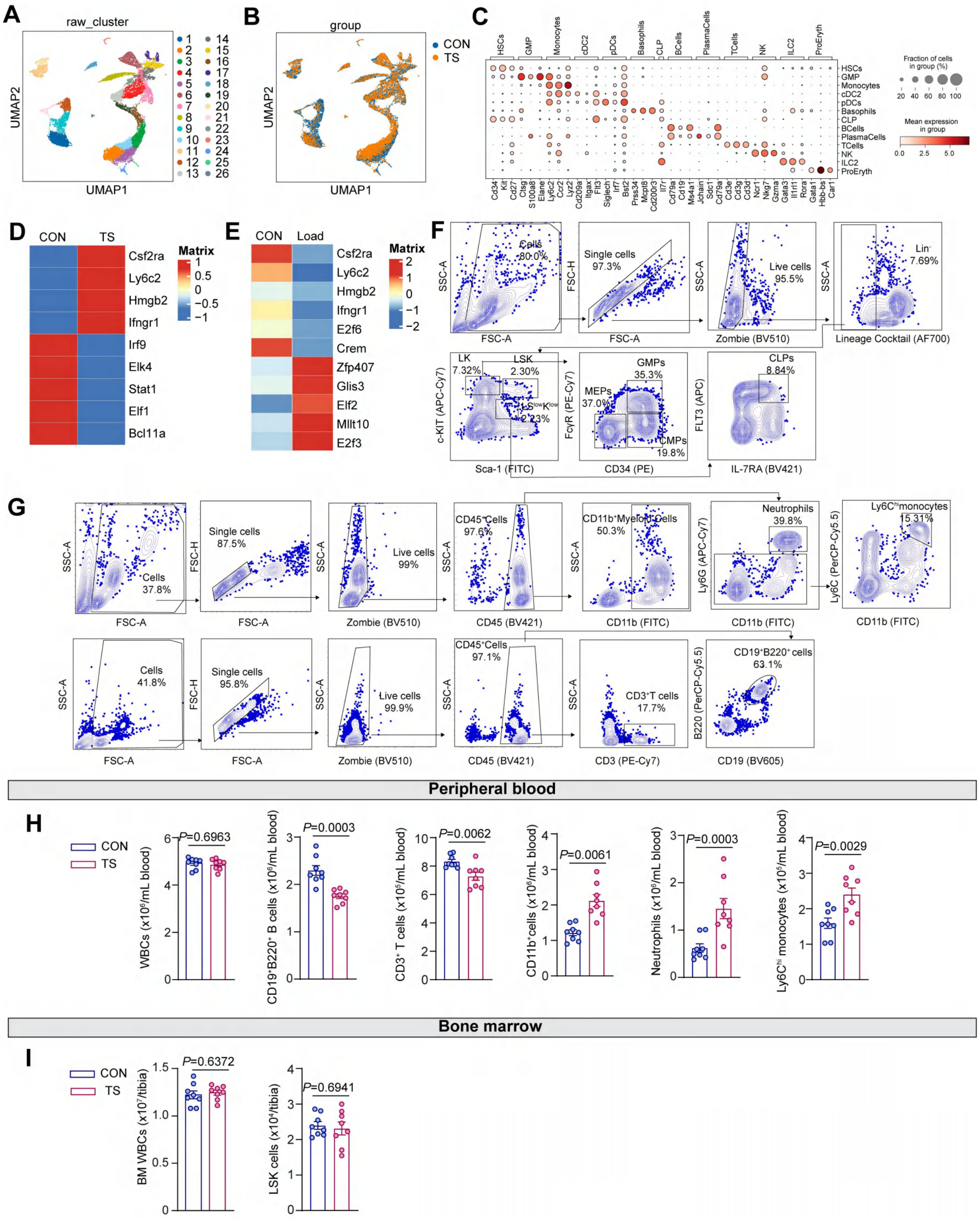


**Figure S1. Single-cell transcriptomic atlas and marker-based annotation of bone marrow haematopoietic populations under mechanical unloading and loading.**

**A,** UMAP visualization of bone marrow cells profiled by scRNA-seq, colored by unsupervised clusters. **B,** UMAP visualization of the same cells colored by experimental group, CON or TS. **C,** Dot plot showing canonical marker genes used for cell-type annotation. **D,** Heatmap showing selected lineage-associated genes in bone marrow progenitor populations from CON and TS mice. **E,** Heatmap showing selected lineage-associated genes in bone marrow progenitor populations from CON and mechanically loaded mice, highlighting reciprocal changes after mechanical loading. **F,** Representative gating strategy for bone marrow hematopoietic stem and progenitor populations. **G,** Representative gating strategy for peripheral blood leukocyte subsets. **H,** Absolute counts of peripheral blood white blood cells and indicated immune subsets in CON and TS mice, including total WBCs, B cells, T cells, CD11b$^{+}$ myeloid cells, neutrophils and Ly6C$^{hi}$ monocytes ($n$ = 8). **I,** Absolute numbers of bone marrow white blood cells and LSK cells in CON and TS mice ($n$ = 8). For scRNA-seq analyses in **A**–**E**, $n$ = 2 mice per group, with each mouse processed as an independent biological replicate. Data are presented as mean ± s.e.m. $P$ values were calculated using two-sided unpaired Student's t-test and are indicated in the plots.

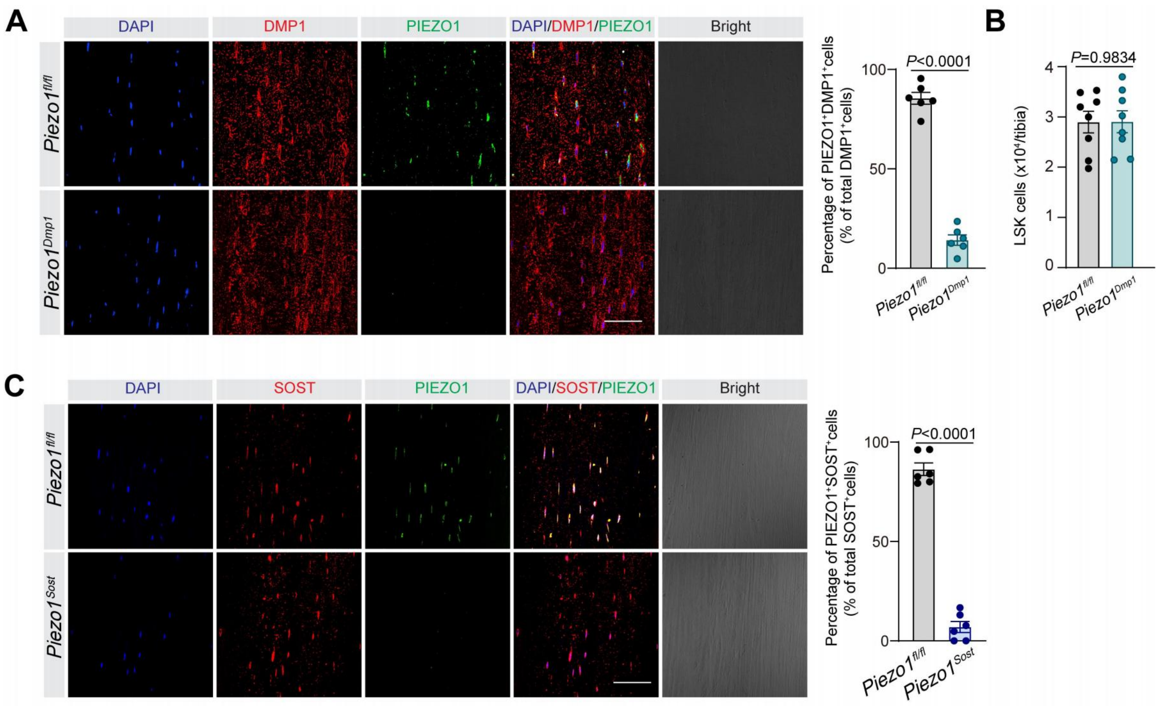


**Figure S2. Validation of osteocyte-lineage *Piezo1* deletion in *Dmp1*-Cre and *Sost*-CreERT2 models.**

**A,** Representative immunofluorescence images of tibial bone sections from *Piezo1*$^{\mathrm{fl/fl}}$ and *Piezo1*$^{Dmp1}$ mice stained for DAPI, DMP1 and PIEZO1. Merged fluorescence images and corresponding bright-field images are shown. Right, quantification of PIEZO1$^+$DMP1$^+$ cells as a percentage of total DMP1$^+$ cells ($n$ = 8). Scale bar, 100 µm. **B,** Absolute numbers of LSK cells in bone marrow from *Piezo1*$^{\mathrm{fl/fl}}$ and *Piezo1*$^{Dmp1}$ mice ($n$ = 8). **C,** Representative immunofluorescence images of tibial bone sections from tamoxifen-treated *Piezo1*$^{\mathrm{fl/fl}}$ and *Piezo1*$^{Sost}$ mice stained for DAPI, SOST and PIEZO1. Merged fluorescence images and corresponding bright-field images are shown. Right, quantification of PIEZO1$^+$SOST$^+$ cells as a percentage of total SOST$^+$ cells ($n$ = 6). Scale bar, 100 µm. Each dot represents one mouse. Data are presented as mean ± s.e.m. *P* values were calculated using two-sided unpaired Student's *t*-test and are indicated in the plots.

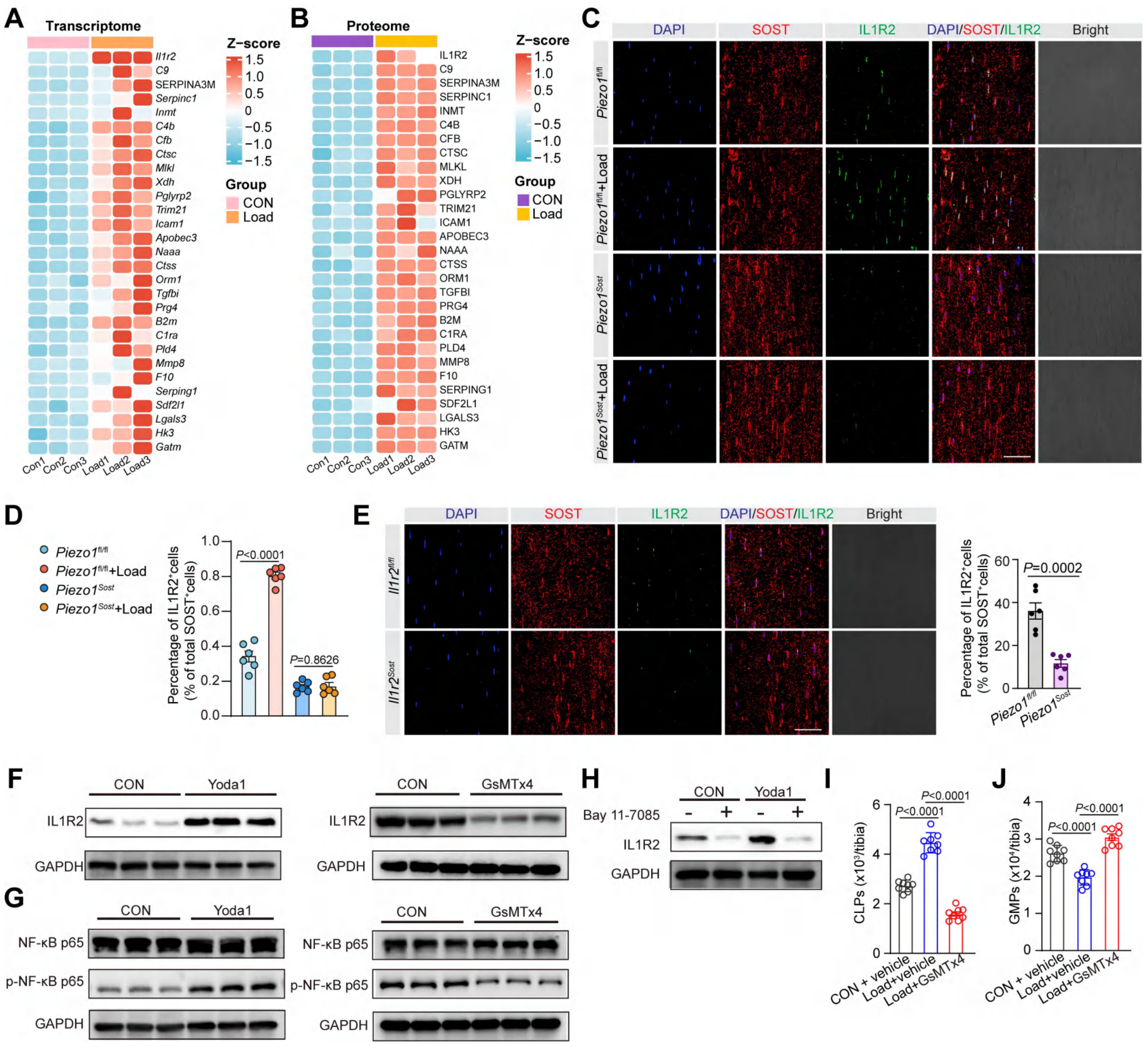


**Figure S3. PIEZO1–NF-κB signaling in osteocytes drives IL1R2 induction and is required for loading-mediated pro-lymphoid hematopoiesis.**

**A,** Heat map (Z-score) of mechanically responsive genes identified from bone transcriptomic profiling (CON vs DTAL; biological replicates shown). **B,** Heat map (Z-score) of differentially abundant proteins identified by quantitative proteomic analysis of BM supernatants from control and DTAL-treated mice (CON vs Load). **C,** Representative immunofluorescence staining of bone sections showing DAPI (blue), SOST (red) and IL1R2 (green) in *Piezo1*$^{fl/fl}$ and *Piezo1*$^{Sost}$ mice with or without DTAL; merged and bright-field images are shown, scale bar, 100 μm. **D,** Quantification of IL1R2 signal in bone (as indicated) demonstrating DTAL-induced IL1R2 upregulation in *Piezo1*$^{fl/fl}$ but not *Piezo1*$^{Sost}$ mice, *n* = 6. **E,** Representative immunofluorescence images of IL1R2 in SOST$^{+}$ osteocytes from *Il1r2*$^{fl/fl}$ and *Il1r2*$^{Sost}$ mice; right, quantification of IL1R2$^{+}$ cells among SOST$^{+}$ osteocytes ($n$ = 8), scale bar, 100 μm. **F,** Immunoblot analysis of IL1R2 in osteocytes following PIEZO1 activation by Yoda1 (left) or inhibition by GsMTx4 (right). **G,** Immunoblot analysis of total NF-κB p65 and phosphorylated NF-κB p65 (p–NF-κB p65) in osteocytes following Yoda1 stimulation (left) or GsMTx4 treatment (right). **H,** Bay 11-7085–mediated NF-κB inhibition blocks Yoda1-induced IL1R2 upregulation in osteocytes, assessed by immunoblot. **I, J,** Pharmacological inhibition of PIEZO1 with GsMTx4 abolishes DTAL-induced expansion of CLPs (**I**) and the reduction of GMPs (**J**), quantified by flow cytometry ($n$ = 8). Data are presented as mean ± s.e.m. For **D**, statistical significance was assessed using two-way ANOVA followed by multiple-comparisons correction. For **E**, two-sided unpaired Student's *t*-test was used. For **I** and **J**, one-way ANOVA followed by multiple-comparisons correction was used. Representative immunoblots from three independent experiments are shown, with GAPDH used as a loading control. Exact *P* values are indicated in the plots.

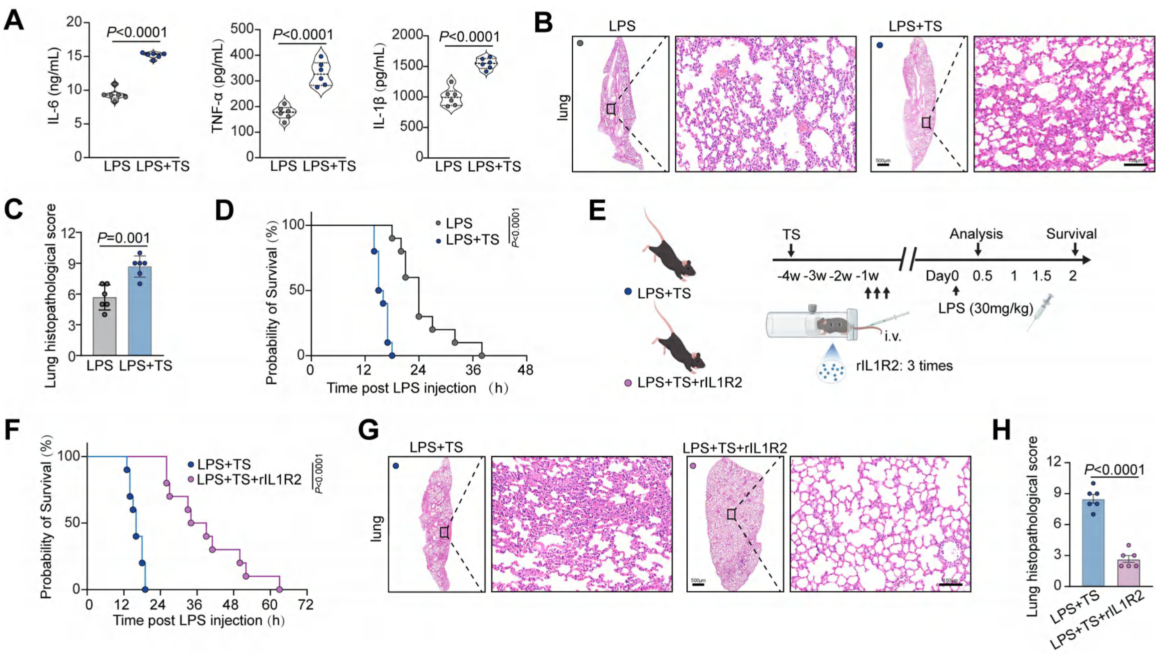


**Figure S4. IL1R2 mitigates mechanical unloading-exacerbated systemic inflammation and lung injury after LPS challenge.**

**A,** Serum concentrations of IL-6, TNF-α and IL-1β after LPS challenge in control mice and mice subjected to prior TS ($n$ = 6). **B,** Representative H&E-stained lung sections from LPS and LPS+TS mice. Boxed regions are shown at higher magnification. Scale bars, 500 μm for overview images and 100 μm for magnified images. **C,** Quantification of lung histopathological scores in LPS and LPS+TS mice ($n$ = 6). **D,** Kaplan–Meier survival curves after LPS challenge in LPS and LPS+TS mice ($n$ = 10). **E,** Experimental scheme for recombinant soluble IL1R2 administration in tail-suspended mice undergoing LPS challenge. Mice were subjected to TS for 4 weeks and then challenged with LPS; recombinant soluble IL1R2 was administered intravenously as indicated. **F,** Kaplan–Meier survival curves of LPS+TS mice treated with vehicle or recombinant soluble IL1R2 (rIL1R2) ($n$ = 10). **G,** Representative H&E-stained lung sections from LPS+TS mice treated with vehicle or rIL1R2. Boxed regions are shown at higher magnification. Scale bars, 500 μm for overview images and 100 μm for magnified images. **H,** Quantification of lung histopathological scores in LPS+TS mice treated with vehicle or rIL1R2 ($n$ = 6). For cytokine measurements in **A**, data are presented as mean ± s.e.m., and $P$ values were calculated using two-sided unpaired Student's $t$-test. For lung histopathological scores in **C** and **H**, data are presented as mean ± s.e.m., and $P$ values were calculated using the Mann–Whitney U test. For survival analyses in **D** and **F**, $P$ values were calculated using the log-rank Mantel–Cox test. Each dot represents one mouse. Exact $P$ values are indicated in the plots.

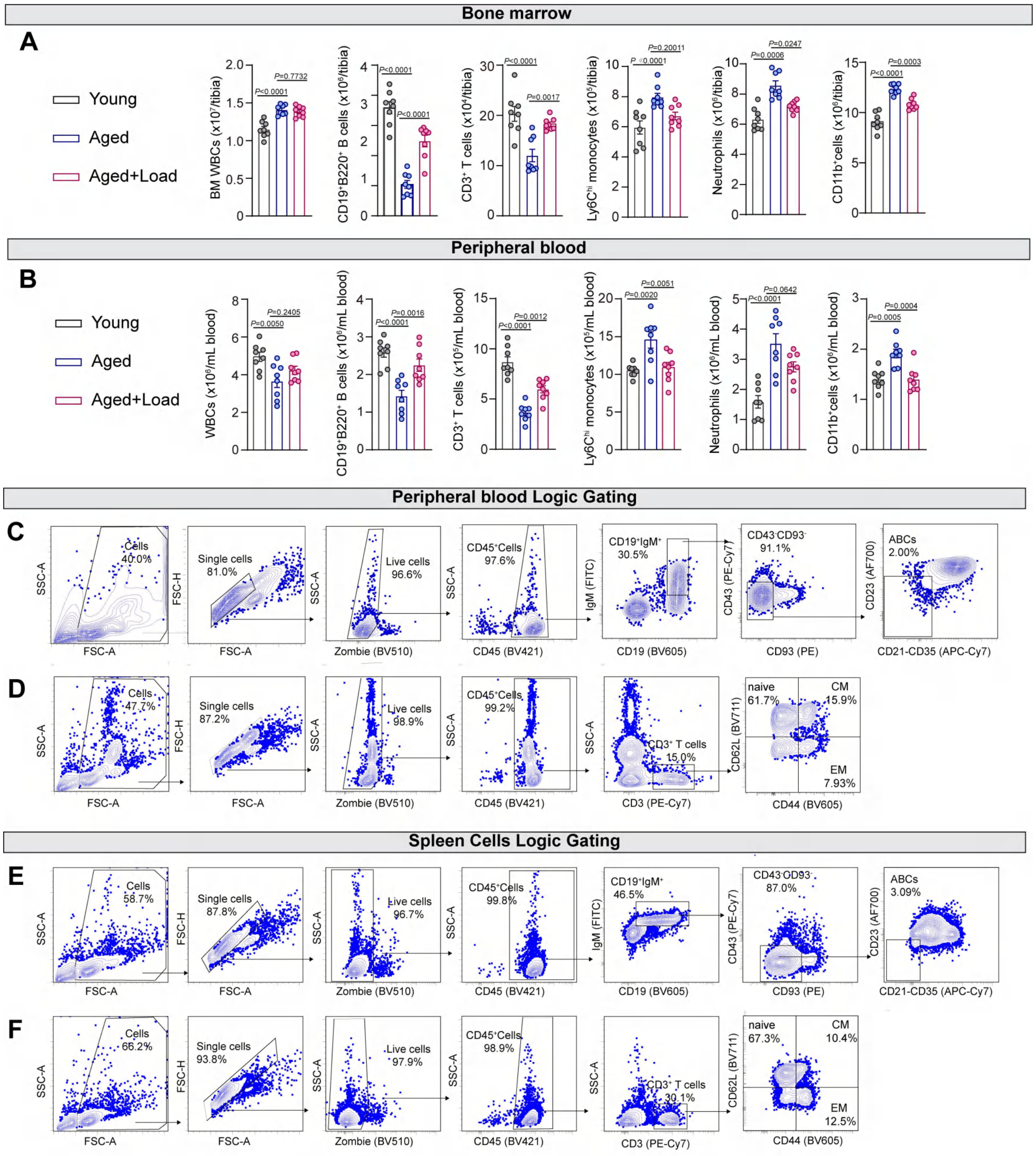


**Figure S5. Mechanical loading mitigates age-associated immune remodeling across bone marrow, blood and spleen.**

**A,** Flow-cytometric quantification of major immune populations in bone marrow (tibia) from young, aged, and aged + DTAL (Aged + Load) mice, including total BM WBCs, $CD19^+$ $B220^+$ B cells, $CD3^+$ T cells, $Ly6C^{hi}$ monocytes, neutrophils, and $CD11b^+$ myeloid cells ($n$ = 8). **B,** Peripheral blood leukocyte profiling from young, aged, and Aged + Load mice, including WBCs, $CD19^+$ $B220^+$ B cells, $CD3^+$ T cells, $Ly6C^{hi}$

monocytes, neutrophils, and CD11b$^{+}$ myeloid cells (normalized to blood volume, as indicated) ($n$ = 8). **C, D,** Representative logical gating for peripheral blood leukocytes. Cells were sequentially gated as Cells → singlets → live cells → CD45$^{+}$ leukocytes. B cells were identified as CD19$^{+}$IgM$^{+}$, and mature B cells were further gated as CD43$^{-}$CD93$^{-}$; age-associated B cells (ABCs) were defined within this population based on CD23 and CD21/CD35 expression (as indicated). T cells were identified as CD3$^{+}$, then separated into CD4$^{+}$ and CD8$^{+}$ subsets. Naive, central memory (CM), and effector memory (EM) T cells were defined by CD62L and CD44 expression (naive: CD44$^{lo}$CD62L$^{hi}$; CM: CD44$^{hi}$CD62L$^{hi}$; EM: CD44$^{hi}$CD62L$^{lo}$). **E, F,** Representative logical gating for spleen leukocytes using the same sequential strategy and marker definitions as in peripheral blood, including identification of CD19$^{+}$IgM$^{+}$ B cells, CD43$^{-}$CD93$^{-}$ mature B cells, ABCs (CD23/CD21-CD35 gate), and CD3$^{+}$ T cells with CD4$^{+}$/CD8$^{+}$ and naive/CM/EM subdivision by CD44/CD62L. Data are presented as mean ± s.e.m. Comparisons among young, aged and aged+Load groups were performed using one-way ANOVA followed by multiple-comparisons correction. Exact $P$ values are indicated in the plots.

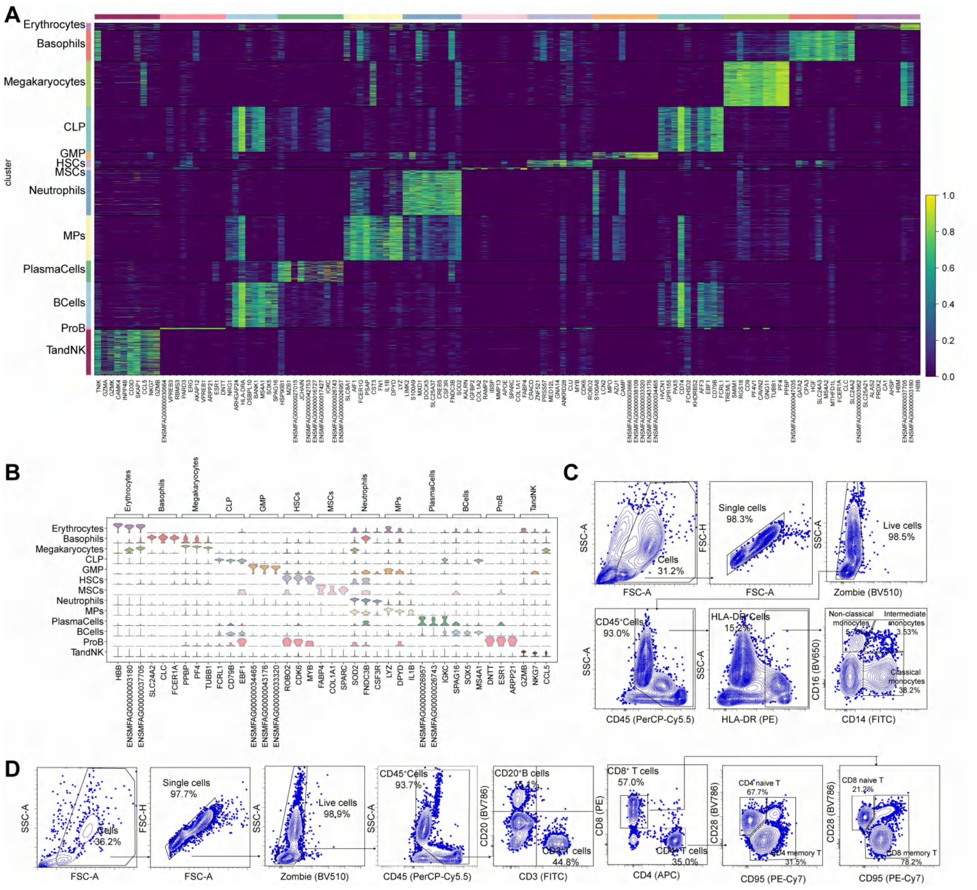


**Figure S6. Marker-based annotation of cynomolgus macaque bone marrow cells and peripheral blood flow-cytometry gating strategies.**

**A,** Heatmap showing scaled expression of representative marker genes across annotated bone marrow cell populations identified by scRNA-seq, including erythrocytes, basophils, megakaryocytes, CLPs, GMPs, HSCs, MSCs, neutrophils, monocytes/macrophages, plasma cells, B cells, pro-B cells and T/NK cells. Cells are grouped by annotated cluster identity, and genes are ordered by lineage-associated expression patterns. **B,** Dot plot showing canonical marker genes used for cell-type annotation across the indicated bone marrow clusters. Dot size indicates the fraction of cells expressing each gene, and color indicates average normalized expression within each cluster. **C,** Representative flow-cytometry gating strategy for peripheral blood monocyte subsets. After exclusion of debris, doublets and dead cells, CD45$^{+}$ leukocytes were gated, followed by identification of HLA-DR$^{+}$ cells. Monocyte subsets were defined based on CD14 and CD16 expression, including classical, intermediate and non-classical monocytes. **D,** Representative flow-cytometry gating strategy for peripheral blood lymphocyte subsets. After exclusion of debris, doublets and dead cells, CD45$^{+}$ leukocytes were gated, followed by identification of CD20$^{+}$ B cells and CD3$^{+}$ T cells. CD3$^{+}$ T cells were further separated into CD4$^{+}$ and CD8$^{+}$ T cells, and naive T cells were defined based on CD28 and CD95 expression.

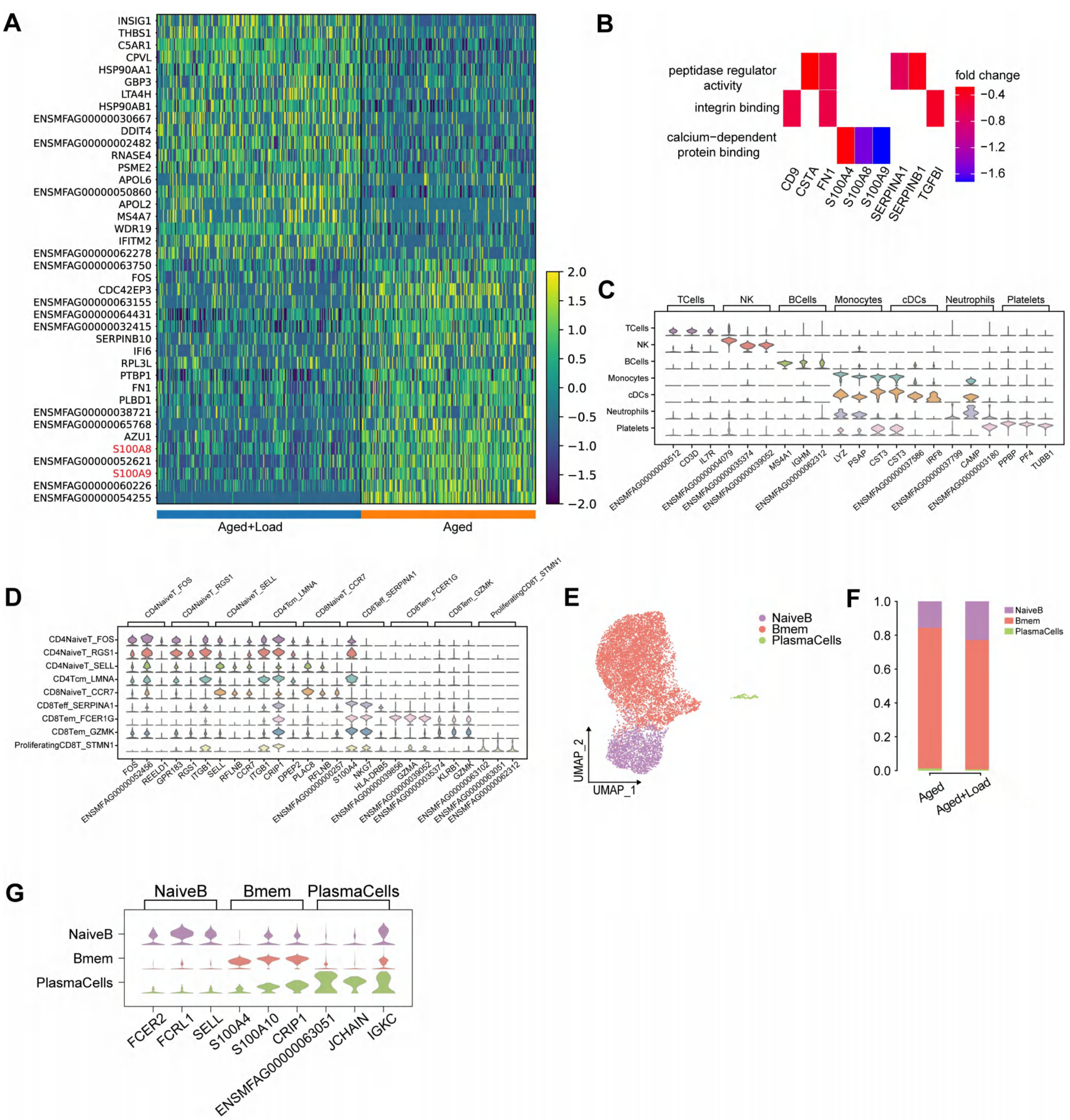


**Figure S7. Mechanical loading reshapes transcriptional programs of peripheral immune cells in**

**aged cynomolgus macaques and alters B-cell composition.**

**A,** Heatmap showing representative differentially expressed genes in PBMCs from aged sham-loaded and aged+Load macaques. Rows represent genes and columns represent single cells; color indicates scaled normalized expression. Inflammation- and myeloid-associated genes, including S100A8 and S100A9, are highlighted in red. **B,** Gene Ontology molecular-function enrichment of genes reduced in aged+Load macaques relative to aged sham-loaded controls. Columns indicate representative contributing genes, and color indicates fold change in aged+Load relative to aged sham-loaded controls. **C,** Violin plots showing canonical marker genes used to annotate major circulating immune lineages, including T cells, NK cells, B cells, monocytes, conventional dendritic cells, neutrophils and platelets. **D,** Violin plots showing representative marker genes used to annotate T-cell states, including $CD4^+$ naive-like, $CD4^+$ central memory-like, $CD8^+$ naive-like, $CD8^+$ effector-like, $CD8^+$ effector memory-like and proliferating $CD8^+$ T-cell states. **E,** UMAP visualization of reclustered B-lineage cells, annotated as naive B cells, memory B cells and plasma cells. **F,** Relative proportions of naive B cells, memory B cells and plasma cells among B-lineage cells in aged sham-loaded and aged+Load macaques. **G,** Violin plots showing representative B-cell subset markers, including naive B-cell markers FCER2, FCRL1 and SELL; memory B-cell-associated markers S100A4, S100A10 and CRIP1; and plasma-cell-associated markers JCHAIN and IGKC. For PBMC scRNA-seq analyses, $n$ = 3 biologically independent macaques per group.
